# A stress-responsive p38 signaling axis in choanoflagellates

**DOI:** 10.1101/2022.08.26.505350

**Authors:** Florentine U. Rutaganira, Maxwell C. Coyle, Maria H.T. Nguyen, Iliana Hernandez, Alex P. Scopton, Arvin C. Dar, Nicole King

## Abstract

Animal kinases regulate cellular responses to environmental stimuli, including cell differentiation, migration, survival, and response to stress, but the ancestry of these functions is poorly understood. Choanoflagellates, the closest living relatives of animals, encode homologs of diverse animal kinases and have emerged as model organisms for reconstructing animal origins. However, efforts to identify key kinase regulators in choanoflagellates have been constrained by the limitations of currently available genetic tools. Here, we report on a framework that combines small molecule-driven kinase discovery with targeted genetics to reveal kinase function in choanoflagellates. To study the physiological roles of choanoflagellate kinases, we established two high-throughput platforms to screen the model choanoflagellate *Salpingoeca rosetta* with a curated library of human kinase inhibitors. We identified 95 diverse kinase inhibitors that disrupt *S. rosetta* cell proliferation. By focusing on one inhibitor, sorafenib, we identified a p38 kinase as a regulator of the heat shock response in *S. rosetta*. This finding reveals a conserved p38 function between choanoflagellates, animals, and fungi. Moreover, this study demonstrates that existing kinase inhibitors can serve as powerful tools to examine the ancestral roles of kinases that regulate modern animal development.

## Introduction

Phenotypic screens with libraries of small molecules have revolutionized cell biology by providing chemical tools to study protein function ^1–4^. Because aberrant kinase activity can lead to human disease ^5,6^, many tools have been developed to inhibit kinase activity and detect protein phosphorylation. Small molecules that target the kinase active site coupled with assays of kinase inhibition have resulted in effective therapeutic strategies to counter the functions of misregulated kinases, including inhibition of aberrant cell growth and proliferation caused by oncogenic kinases ^5,7^.

We sought to test whether kinase-regulated physiology in choanoflagellates, the closest living relatives of animals, could be revealed by kinase inhibitors. Choanoflagellates possess homologs of diverse animal kinases (Figure S1) ^8–10^ and, due to their phylogenetic placement, are well-suited for studies of the ancestral functions of animal cell signaling proteins ^11,12^. Indeed, a previous study showed that two broad-spectrum kinase inhibitors disrupt cell proliferation in the choanoflagellate *Monosiga brevicollis* ^13^. However, this study did not demonstrate whether kinases were directly targeted or identify specific pathways regulated by kinase signaling.

Using a library of well-characterized kinase inhibitors that vary in their human kinase inhibition profile, we treated cultures of *Salpingoeca rosetta*, a model choanoflagellate, in a multiwell format. We found that treatment of *S. rosetta* cultures with a set of kinase inhibitors disrupted cell proliferation and led to global inhibition of *S. rosetta* phosphotyrosine signaling. Using one of these inhibitors, sorafenib, followed by reverse genetics, we found that an *S. rosetta* p38 kinase homolog is activated by environmental stressors and signals downstream of sorafenib-inhibited kinases.

## Results

### Screening of a human kinase inhibitor library reveals small molecules that inhibit *S. rosetta* kinase signaling and cell proliferation

To investigate whether kinase activity regulates *S. rosetta* cell proliferation, we treated *S. rosetta* cultures with characterized human kinase inhibitors. Because the highest conservation between choanoflagellate and human kinases occurs in the kinase domain ^8–10^, we focused on kinase inhibitors that bind in the kinase active site. As a proof-of-concept, we first assayed staurosporine, a well-characterized broad-spectrum kinase inhibitor and inducer of cell death in diverse organisms ^14,15^. Our initial screen used flow cytometry to measure the density of *S. rosetta* cells individually treated with staurosporine in multiwell plates. Staurosporine significantly and reproducibly reduced *S. rosetta* cell density (Figure S2A-C) and tyrosine phosphorylation (Figure S2D) in a dose-dependent manner.

After validating the flow cytometry pipeline with staurosporine, we expanded our study to screen 1255 inhibitors of diverse human kinases (Figure S1, Table S1). We screened *S. rosetta* cultures with each of the molecules in the library at 10 µM and measured cell density at a 24-hour endpoint (Figure 1A and B, Table S1). This screen revealed 44 compounds (3.5% of the library; Figure 1A and Figure S3) that significantly decreased *S. rosetta* cell counts compared to DMSO controls.

**Figure 1.**
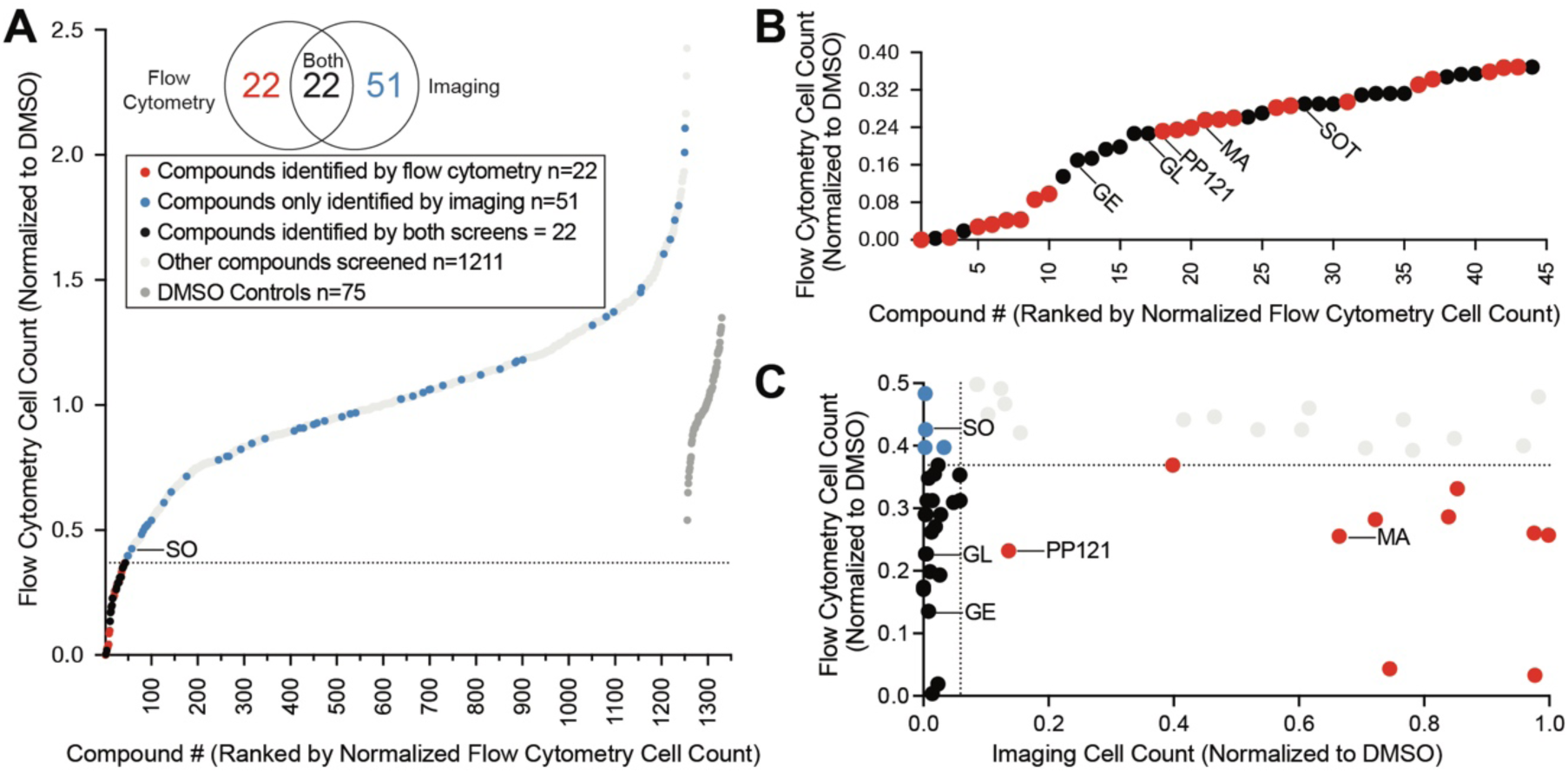
High-throughput screening of a small molecule library revealed inhibitors of *S. rosetta* cell proliferation. (A) Treatment of *S. rosetta* cultures with 1255 different small molecules (see Table S1) resulted in a distribution of cell counts, assessed by flow cytometry, at the 24-hour endpoint. *S. rosetta* cell counts were normalized to the average of DMSO controls within the same plate (dark grey). Compounds determined to significantly inhibit *S. rosetta* cell proliferation (based on two-tailed p-value < 0.05 calculated from z-score), fall below the dotted line and are indicated in red. Compounds that were not detected as significant inhibitors by flow cytometry but were identified by imaging (based on two-tailed p-value < 0.05 calculated from z-score) are in blue. Compounds that were not significant inhibitors for either screen are indicated in light grey. Sorafenib (SO), a focus of this study, is labeled. (B) The range of normalized cell counts measured by flow cytometry for compounds that significantly inhibited *S. rosetta* cell proliferation. Compounds that were the focus of further study – genistein (GE), glesatinib (GL), PP121, masitinib (MA), sotrastaurin (SOT) – are labeled. (C) Comparison of normalized values of compounds that inhibited *S. rosetta* cell proliferation, assessed by flow cytometry and the corresponding normalized values determined by imaging. Compounds determined to significantly inhibit *S.* rosetta cell proliferation (based on two-tailed p-value < 0.05 calculated from z-score) by flow cytometry fall below the dotted line on the y-axis and by imaging, to the left of the dotted line on the x-axis.

As a complementary assay, we pursued an imaging-based workflow to measure cell density. Although *S. rosetta* cell density could be measured by flow cytometry without staining reagents, our co-culture system and treatment paradigm presented two potential sources of inaccuracy: compound aggregation due to low solubility in choanoflagellate media and choanoflagellate-sized clumps of bacterial biofilm. Therefore, we developed an imaging pipeline to enumerate *S. rosetta* cells by segmenting fixed-cell immunofluorescence micrographs at a 48-hour endpoint. Our imaging pipeline distinguished wells with staurosporine-treated cells from DMSO controls (Figure S2E, F) with comparable z’ standard statistics to flow cytometry (Figure S2B, S2E) ^16^. This orthogonal approach identified 22 compounds (1.8% of the library) that overlapped with the flow cytometry screen and 51 additional molecules (4.4% of the library) that inhibited *S. rosetta* cell proliferation but were not identified in our flow cytometry screen (Figure 1A, Figure 1C, Figure S2G, Figure S3). In total, 95 ATP-competitive inhibitors of human protein and lipid kinases (Figure S4A-C) that ranged in selectivity (Figure S4D) were identified as potential inhibitors of *S. rosetta* cell proliferation.

Choanoflagellates are predicted to express kinases that regulate animal cell growth, including mitotic kinases ^17^, the serine-threonine kinase Akt ^18^, and a diverse set of tyrosine kinases ^8^. We identified GSK461364 and Volasertib, inhibitors of polo-like kinase 1 (PLK1), a mitotic kinase, in both screens (Figure S3). In addition, both screens identified diverse inhibitors of human Akt and tyrosine kinases (Figure S3). We also identified inhibitors of *S. rosetta* cell proliferation that disrupt human kinase signaling indirectly, i.e. without binding to a kinase (Figure S3). For example, genistein, a natural product that indirectly alters kinase activity ^19,20^, inhibited *S. rosetta* cell proliferation and was previously shown to inhibit the growth of *M. brevicollis* ^13^.

### Some inhibitors of S. rosetta cell proliferation also disrupt S. rosetta phosphotyrosine signaling

After identifying human tyrosine kinase inhibitors that inhibited *S. rosetta* cell proliferation, we used biochemical methods to determine if the phenotype observed was correlated with inhibition of *S. rosetta* kinase activity. Widely available phospho-specific antibodies that distinguish phosphorylated amino acids from their unphosphorylated cognates can reveal kinase activity in diverse organisms ^13,21–24^ and have been used to detect tyrosine phosphorylation of peptides or animal proteins by heterologously expressed choanoflagellate kinases ^25–28^. We triaged our set of 95 identified proliferation inhibitors to focus on those compounds that inhibit human tyrosine kinases (Figure S3), as opposed to those that inhibit serine or threonine phosphorylation, in part because of the relative lack of specificity of commercially-available phosphoserine and phosphothreonine antibodies ^22–24^.

In particular, we focused on four inhibitors – sorafenib, glesatinib, masitinib and PP121 – that were identified by either or both screening paradigms (Table S1, Figure S3) and have a narrower range of kinase targets than staurosporine ^29–32^. In humans, sorafenib, glesatinib, masitinib and PP121 inhibit select receptor tyrosine kinases (RTKs), among other targets. Sorafenib also inhibits some non-receptor tyrosine kinases, tyrosine kinase-like serine-threonine kinases, and p38 stress-responsive kinases ^30^. Masitinib and PP121 also inhibit Src kinase and PP121 additionally inhibits PI3K ^29–31^. Specific residues shared in the kinase domains of human and choanoflagellate tyrosine kinases suggested that these inhibitors might effectively inhibit choanoflagellate tyrosine kinases. For example, choanoflagellate tyrosine kinases have residues within the predicted active site that are necessary for kinase activity in other organisms (e.g. “K” in VAIK, “D” in HRD and DFG) (Figure S5A) ^8,33–35^ and additional residues that confer sensitivity towards tyrosine kinase inhibitors (Figure S5B-D) ^36,37^.

Of the four inhibitors tested, *S. rosetta* cultures were most sensitive to sorafenib and glesatinib. Although 1 µM masitinib and PP121 were sufficient to reduce *S. rosetta* cell proliferation over the first 40 hours of treatment (Figure 2A), masitinib- and PP121-treated cultures recovered within 85 hours. Masitinib and PP121 are effective in human culture media for 48-72 hours ^31,38^, so we have no reason to believe this recovery is due to reduced inhibitor stability, although we can’t rule it out. Of note, both compounds were found to be ineffective inhibitors of cell proliferation in the imaging pipeline, which had an endpoint of 48 hours (Figure 1C).

**Figure 2.**
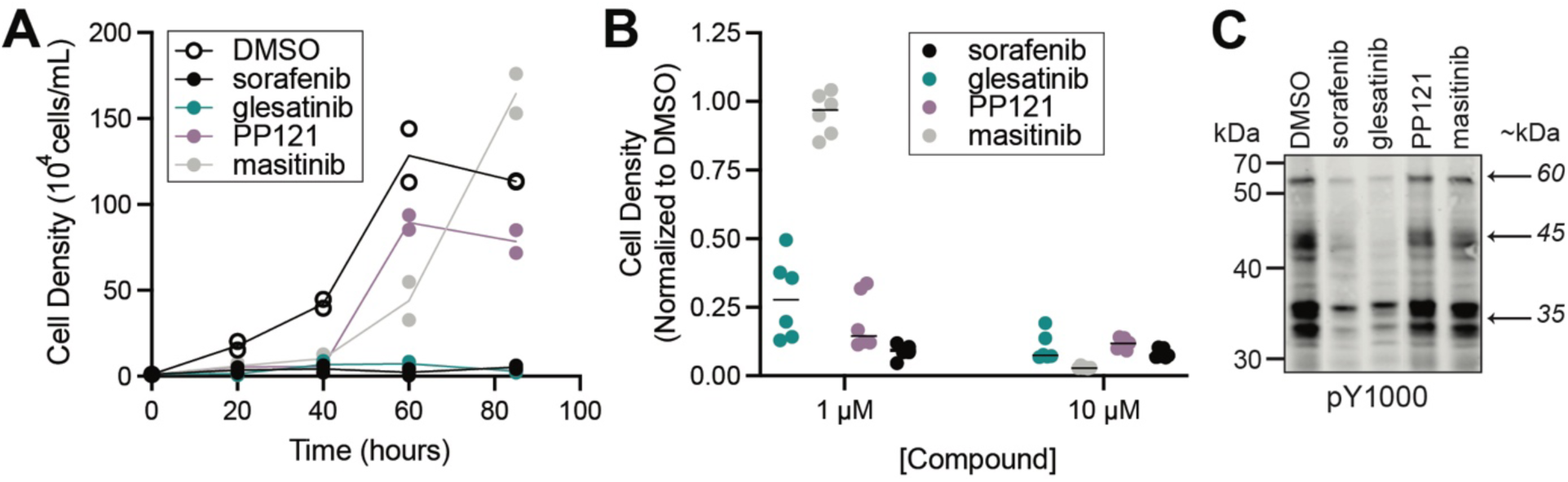
Glesatinib and sorafenib, two multi-target tyrosine kinase inhibitors, disrupt *S. rosetta* cell proliferation and tyrosine phosphosignaling. (A) Treatment of *S. rosetta* cultures with 1 µM sorafenib and glesatinib led to a complete block of cell proliferation, while treatment with 1 µM masitinib or PP121 led to a partial reduction in cell proliferation relative to DMSO-treated cultures. Two biological replicates were conducted per treatment, and each point represents the mean of three measurements from each biological replicate. For timepoints at 40, 60, and 85 hours, cell densities of inhibitor-treated cultures were significantly different from vehicle (DMSO) (p-value <0.01). Significance was determined by a two-way ANOVA multiple comparisons test. (B) *S. rosetta* cultures treated with 1 µM or 10 µM sorafenib, glesatinib, or PP121 for 24 hours had reduced normalized cell density, whereas masitinib only had reduced normalized cell density at 10 µM. Normalized cell densities were determined to be reduced if differences between treatments and vehicle (DMSO) were significant (p-value <0.01) Significance was determined by determined by a two-way ANOVA multiple comparisons test. Movies show *S. rosetta* cells treated with 10 µM glesatinib that undergo cell lysis (Movie S1) and sorafenib, that have cell body deformation (Movies S2-S3), in comparison to DMSO control (Movie S4). (C) Western blot analysis of *S. rosetta* cultures treated with 1 µM sorafenib and glesatinib for 1 hour showed a decrease in tyrosine phosphorylation of proteins at ∼60kDa, ∼45kDa, and ∼35kDa (indicated by arrows and detected with pY1000 anti-phosphotyrosine antibody) compared to vehicle (DMSO) control. Masitinib and PP121 did not reduce the phosphotyrosine signal.

In contrast, sorafenib and glesatinib inhibited cell proliferation throughout the 85-hour growth experiment at 1 µM (Figure 2A) and decreased cell density at 1 µM and 10 µM (Figure 2B). Glesatinib treatment induced cell lysis (Movie S1), whereas sorafenib induced cell body elongation (Movies S2-S3), in comparison to the DMSO control (Movie S4). Importantly, treatment of *S. rosetta* cultures with 1 µM sorafenib or glesatinib led to a global decrease in phosphotyrosine signal, while treatment with 1 µM masitinib or PP121 did not decrease phosphotyrosine levels as detected by western blot (Figure 2C).

These findings showed that some kinase inhibitors, including sorafenib and glesatinib, could disrupt both *S. rosetta* cell proliferation and tyrosine kinase signaling. To further investigate patterns among human kinase inhibitors that showed this effect and identify kinases that might be relevant for the observed inhibition, we tested a panel of 17 human TK inhibitors (Figure S6) that share overlapping kinase targets with sorafenib and glesatinib. Of the 17 compounds tested, treatment with four additional small molecules (regorafenib, AD80, milciclib, and vemurafenib) led to a global decrease in phosphotyrosine staining (Figure S6A, “*”) but not phosphoserine and phosphothreonine staining (Figure S7) as detected by western blot. These four inhibitors mildly reduced the rate of cell proliferation (Figure S6B, Figure S8). Other tyrosine kinase inhibitors impaired *S. rosetta* cell proliferation but did not disrupt phosphotyrosine signaling, including PP2 (Figure S6C, Figure S8), consistent with prior findings in *M. brevicollis*^13^.

### Sorafenib binds to S. rosetta p38 kinase

To identify specific *S. rosetta* kinases whose activity might regulate *S. rosetta* cell physiology, we focused on sorafenib, an inhibitor that binds TKs and serine-threonine kinases ^30^ with well-characterized structure-activity relationships ^39,40^. We started by using ActivX ATP and ADP probes ^41,42^ to covalently enrich for kinases and high-affinity kinase interactors present within *S. rosetta* lysates after pretreatment with vehicle (DMSO) or sorafenib. While sorafenib is predicted to bind to the active site of a select subset of kinases, ActivX probes target ATP and ADP binding proteins more broadly. Therefore, pretreatment of *S. rosetta* lysates with sorafenib was predicted to competitively inhibit binding of ActivX probes to sorafenib targets. By identifying kinases from *S. rosetta* lysates that were more often bound to ActivX probes after DMSO pretreatment of *S. rosetta* lysate compared with those that were recovered by ActivX probes after sorafenib pretreatment, we aimed to identify likely targets of sorafenib. Through this strategy, we identified a predicted *S. rosetta* p38 kinase (Figure S9A) that bound 10-fold less well to ActivX probe in the presence of sorafenib (Figure 3, Table S2). Hereafter, we refer to this serine-threonine kinase as *Sr*-p38 ^43^.

**Figure 3.**
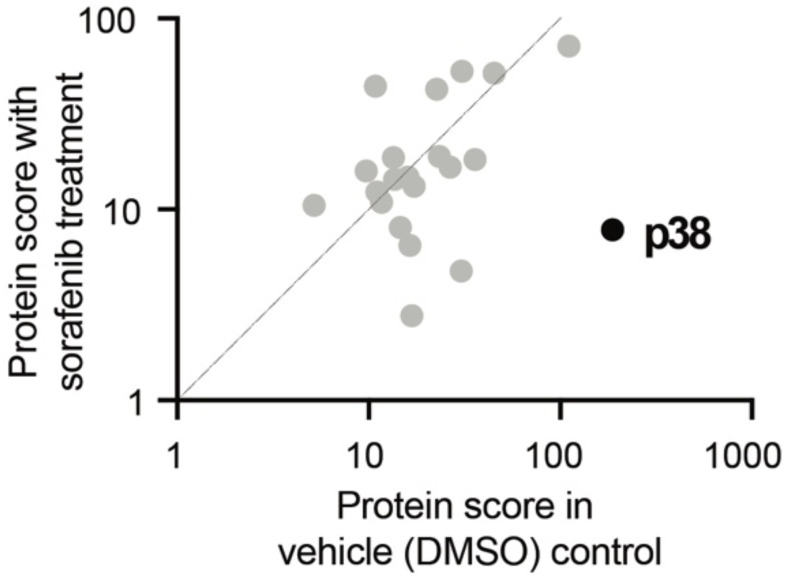
*S. rosetta* p38 binds to Sorafenib. The ActivX ATP probe was used to pull down kinases from *S. rosetta* lysates that were pretreated with either DMSO or the ATP-competitive inhibitor sorafenib. We found that pretreatment with sorafenib reduced the level of p38 recovered using the ActivX ATP probe, indicating that sorafenib and p38 interact and outcompete ActivX ATP probe binding. Kinases plotted are only those that were identified in both vehicle and sorafenib pre-treatments. For full kinase enrichment list, see Table S2, and for alignment of *S. rosetta* p38 with those from animals and fungi, see Fig. S9A.

Kinase enrichment with ActivX probes and other approaches have previously identified human p38 kinases as primary targets of sorafenib ^39,44^. In animals, sorafenib preferentially binds kinases with threonine gatekeeper residues ^39^, including tyrosine kinases and two of four vertebrate p38 kinase paralogs with threonine gatekeepers ^39^ (Figure S9A). *Sr*-p38, contains a threonine gatekeeper (Figure S9A) and lysine residues necessary for ActivX probe binding. Because sorafenib displaced ActivX probe binding to *Sr-*p38, we infer that sorafenib binds to *Sr-* p38 directly. Although sorafenib treatment reduces *S. rosetta* cell proliferation and sorafenib binds to *Sr*-p38, these two findings did not directly implicate *Sr*-p38 in the regulation of cell proliferation. Therefore, we next sought to understand how *Sr-*p38 regulates *S. rosetta* cell physiology.

### S. rosetta p38 is a heat-responsive kinase

Environmental stressors activate p38 kinases in animals and fungi ^43,45–51^ and p38 kinase is present in diverse choanoflagellates (Figure S9B). However, the roles of p38 kinase in choanoflagellate biology are unknown. Although a previous study identified a nutrient-sensitive protein in *M. brevicollis* with a molecular weight similar to p38 kinases ^13^, the identity and function of this protein were not studied directly.

We wondered if stressors relevant to choanoflagellates would activate *Sr-*p38 signaling. Some choanoflagellates, including *S. rosetta*, live in sun-lit water zones that undergo daily and yearly fluctuations in temperature and nutrients ^52–55^. Antibodies specific to phospho-p38 are commercially available, allowing us to detect p38 phosphorylation under different conditions. We found two – p38 MAPK (pThr180/pTyr182) from Biorad and Anti-ACTIVE® p38 from Promega – that recognized a stress-induced protein with minimal background. By generating knockouts of *Sr*-p38 (Methods), we confirmed that the stress-induced protein detected by these antibodies was *Sr*-p38 (Figures 4A-4C). (Unfortunately, the Anti-ACTIVE® p38 antibody from Promega is no longer commercially available, but we provide these results as additional evidence for the connection of *Sr-*p38 to *S. rosetta* cell physiology).

**Figure 4.**
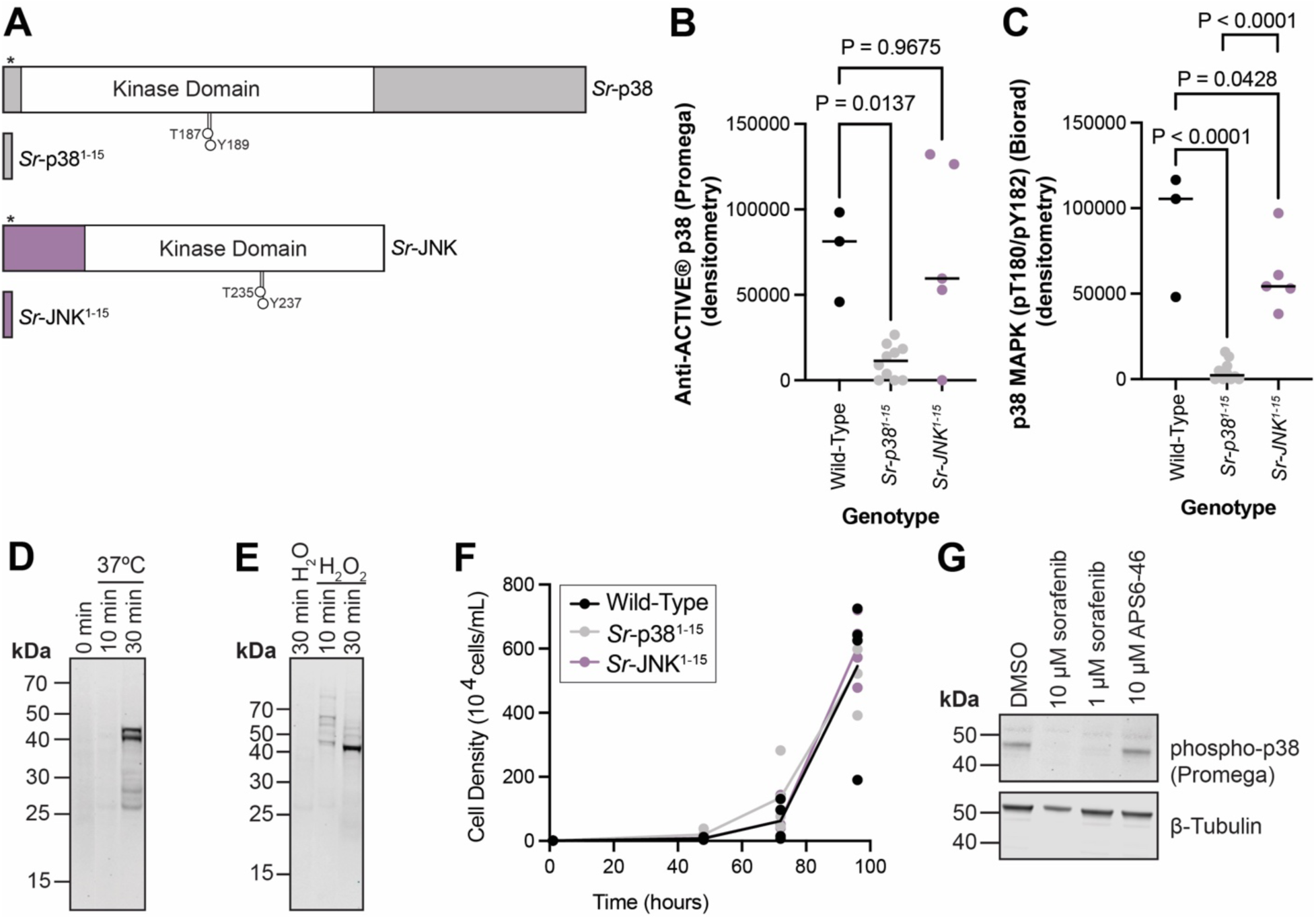
*S. rosetta* p38 phosphorylation is induced by environmental stressors. (A) Strategy for generating *Sr-p38* and *Sr-JNK* knockout cell lines. The *Sr-p38* and *Sr-JNK* loci were targeted by a guide RNA complexed with Cas9 that anneals before the kinase domain and directs Cas9 to introduce a double-strand break downstream of codon 15 (Codon 15 is Serine in Sr-p38 and Alanine in Sr-JNK, indicated by (*). The Cas9-guide RNA complex was coupled with a double-stranded homology-directed repair to introduce a palindromic premature termination stop sequence and a puromycin resistance cassette. The resulting truncated proteins, *Sr*-p38^1–15^ and *Sr*-JNK^1–15^, lack the kinase domain and phosphorylation sites (indicted by the extended circle) recognized by both phospho-p38 antibodies used in this study. Protein diagrams were created with IBS 2.0 ^85^. (B) Heat shock induces Sr-p38 phosphorylation in wild-type cells and the phospho-p38 signal is recognized by the Anti-ACTIVE® p38 antibody (Promega #V1211). This phospho-p38 signal is decreased in *Sr-p38*^1–15^ knockout cell lines but not *Sr-JNK*^1–15^ knockout lines, indicating that the Anti-ACTIVE® p38 antibody (Promega #V1211) detects *Sr-* p38 and that *Sr-*p38, but not *Sr-*JNK, responds to heat shock. Three biological replicates of wild-type cells, ten clones of *Sr-p38*^1–15^ and five clones of *Sr-JNK*^1–15^ strains were incubated at 37°C for one hour. Lysates from the treated cultures were analyzed by western blot with the Anti-ACTIVE® p38 antibody and quantified by densitometry to identify if any changes in *Sr*-p38 phosphorylation occurred. Significance was determined by a one-way ANOVA multiple comparisons test between wild-type cells and *Sr-p38*^1–15^ or *Sr-JNK*^1–15^. (C) Similar to (B), the phospho-p38 signal recognized by the p38 MAPK pThr180/pTyr182 (Biorad #AHP905) antibody in heat shocked wild-type cells is decreased in *Sr-p38*^1–15^ knockout cells but not *Sr-JNK*^1–15^ knockout cells. (D) *S. rosetta* cells, normally cultured at 22°C were incubated at 37°C to induce heat shock. Lysates from the treated cultures were analyzed by western blot with the Anti-ACTIVE® p38 antibody (Promega #V1211) to identify if any changes in *Sr*-p38 phosphorylation occurred. 30 minutes of heat shock was sufficient to induce *Sr*-p38 phosphorylation. (E) *S. rosetta* cells were treated with hydrogen peroxide, a form of oxidative stress for 10 min. and 30 min. 10 min of treatment with 0.5M H2O2 at 22°C was sufficient to induce *Sr*-p38 phosphorylation detected by the Anti-ACTIVE® p38 antibody (Promega #V1211) (F) *Sr-p38*^1–15^ and *Sr-JNK*^1–15^ strains grow similarly to wild-type. Four wild-type cultures and four randomly selected *Sr-p38*^1–15^ and *Sr-JNK*^1–15^ clones were grown in 24-well plates over a 96-hour growth course and showed similar growth. Significance was determined by a two-way ANOVA multiple comparisons test. (G) The induction of *Sr*-p38 phosphorylation by heat shock was kinase-dependent. *S. rosetta* cultures pretreated with 10 µM or 1 µM sorafenib for 30 minutes followed by 30 minutes of heat shock at 37°C and probed with the Anti-ACTIVE® p38 antibody (Promega #V1211) had decreased *Sr*-p38 phosphorylation. APS6-46 treated cultures were not different from vehicle (DMSO) control. In (D), (E), and (G), 4-12% Bis-Tris SDS-PAGE gels were used to resolve the bands observed.

To investigate if *Sr-*p38 is activated in response to environmental stressors, we exposed *S. rosetta* cultures to heat shock and oxidative stress. When *S. rosetta* cells cultured at ambient temperature were subjected to heat and oxidative stress, we observed an increase in phosphorylation (Figures 4B-4E, S10A, S10B). Within 30 minutes of heat shock at 37°C, the phospho-p38 signal increased relative to pre-treatment (Figures 4D, S10A). We also observed an increase in phosphorylation of an ∼45 kDa protein in cell lysates treated with 0.5M hydrogen peroxide (Figure 4E, Figure S10B).

In animals, three stress-activated kinases mediate responses to heat shock and oxidative stress: p38, c-Jun N-terminal kinase (JNK), and extracellular signal-related kinase 5 (ERK5). Upstream dual-specificity kinases (MAP2Ks) activate these kinases through dual phosphorylation of threonine and tyrosine in a short motif: “TGY,” “TPY,” and “TEY” for p38, JNK, and ERK5, respectively ^49,56^ (Figure S9A). Although neither of the phospho-specific p38 antibodies used in this study recognize phosphorylated human JNK or ERK5, the size of the band recognized by these antibodies (∼45 kDa) is closer to the predicted size for *S. rosetta* JNK (39 kDa) than for *S. rosetta* p38 (60 kDa). To test whether *Sr-p38* and *Sr-JNK* was responsive to heat shock and oxidative stress, we generated *Sr-p38* and *Sr-JNK* knockout cell lines using a recently established protocol ^57^ (Figure 4A).

The phospho-p38 signal was reduced in heat-shocked *Sr-p38*^1–15^ lines but not in *Sr-JNK*^1–15^ lines (Figures 4B-4C, S12A-S12B), demonstrating that *Sr-*p38 is the stress-responsive protein detected in our assays. In contrast, the hydrogen peroxide-induced protein phosphorylation signal was preserved in both *Sr-p38*^1–15^ lines and *Sr-JNK*^1–15^ lines (Figures S12D, E). Therefore, heat shock induces phosphorylation of *Sr*-p38, but not *Sr-*JNK. These findings reveal that the connection between heat shock stress and the activation of p38 kinase is conserved between yeast, animals, and choanoflagellates ^43,58,59^.

### Sorafenib inhibits *S. rosetta* cell proliferation separately from the stress-responsive p38 kinase signaling axis

Surprisingly, we found that the growth rates of *Sr-p38*^1–15^ and *Sr-JNK*^1–15^ clones were indistinguishable from those of wild-type cells, indicating that *Sr-*p38 and *Sr-*JNK are dispensable for the regulation of cell proliferation (Figure 4F). Moreover, mutants with reduced sorafenib binding ^39^ maintained sensitivity to sorafenib (Figure S13). Together, these two findings suggest that kinases other than *Sr*-p38 were more relevant to sorafenib’s effect on proliferation.

Previous studies have linked phosphotyrosine signaling to p38 kinase activation in animals in response to multiple stimuli (e.g. heat shock, oxidative stress, growth factors, ultraviolet light) ^49,60^. Sorafenib blocks this signaling axis by inhibiting p38 and upstream kinases (e.g. tyrosine kinases, dual-specificity kinases) ^39,40^. Therefore, we set out to test whether the signaling axis between upstream kinases and p38 in animals is conserved in *S. rosetta*. To this end, we tested whether sorafenib could reduce the observed phosphorylation of *Sr-*p38 in *S. rosetta* cultures subjected to heat shock. As a control, we used APS6-46, a sorafenib analog that shares sorafenib’s core structure but has modifications that make it too large to bind to most sorafenib kinase targets ^61^. Under standard growth conditions, APS6-46 did not inhibit *S. rosetta* tyrosine phosphorylation or cell division (Figure S14), and a previous study found that similar sorafenib analogs could control for off-target inhibition of non-kinase proteins by sorafenib ^61^. In heat-shocked cells, *Sr*-p38 signaling was not activated in *S. rosetta* cultures pretreated with sorafenib, whereas cultures pretreated with APS6-46 (Figure 4G) or selective p38 kinase inhibitors (Figure S15) retained *Sr*-p38 phosphorylation comparable to the DMSO control. Because sorafenib treatment blocked *Sr-*p38 activation and inhibited phosphotyrosine signaling, we infer that sorafenib blocks *Sr*-p38 signal transduction through the inhibition of upstream kinases that transduce the heat stress response in *S. rosetta* (Figure 5). Future studies will be required to identify additional binding targets of sorafenib that regulate *S. rosetta* cell proliferation and heat-responsive activation of *Sr-*p38 (Figure 5).

**Figure 5.**
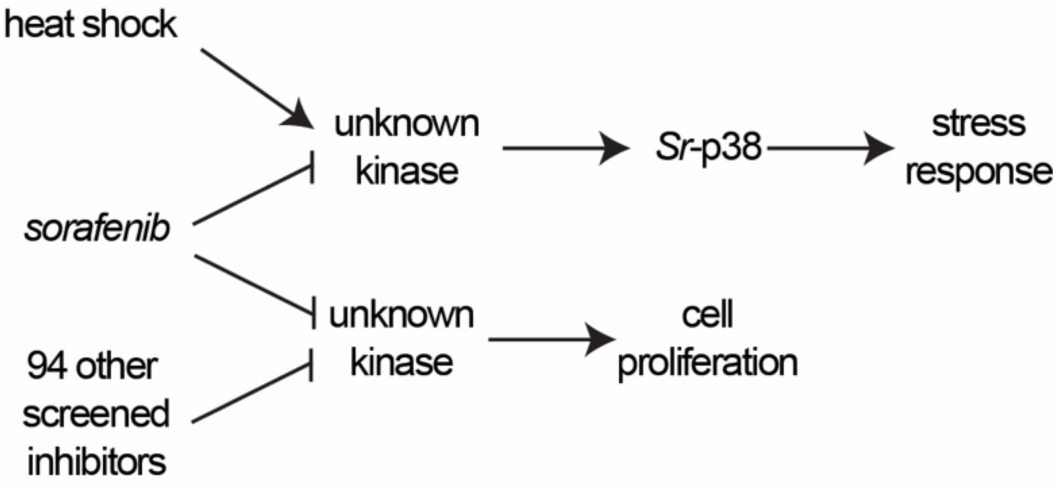
*Sr*-p38 regulates the heat shock response in *S. rosetta*. Proposed mechanism for regulation of the stress-responsive *Sr-*p38 axis. *Sr-*p38 is phosphorylated by upstream kinases in response to heat shock. Sorafenib, a multi-kinase inhibitor, targets kinases upstream of *Sr*-p38 and disrupts *Sr*-p38 signaling. Separately, sorafenib and 94 other small molecules inhibit *S. rosetta* cell proliferation by targeting unknown kinases that regulate *S. rosetta* cell proliferation.

## Discussion

To investigate the relevance of kinase signaling in choanoflagellate cell physiology, we have established two high-throughput phenotypic screens of cells treated with a small molecule library. By treating *S. rosetta* cultures with validated human kinase inhibitors, we uncovered molecules that revealed the physiological relevance of kinases as regulators of *S. rosetta* cell proliferation. Moreover, we identified biologically relevant environmental stressors that activate p38 kinase signaling in *S. rosetta*.

Until recently, the functions of stress-responsive kinases have only been characterized in animals (p38, JNK, and ERK5)^49,56^ and fungi (Hog 1, a homolog of p38) ^43,58,59^. Our discovery that *S. rosetta* phosphorylates p38 and other proteins in response to heat and oxidative shock (Figures 4B-4E) demonstrates that choanoflagellates undergo stress-responsive signaling. Because the phosphorylation of p38 kinases can be inhibited by sorafenib (Figure 4G), we infer that *Sr*-p38 functions within a signaling axis downstream of sorafenib-targeted kinases. Further exploration of this heat-responsive pathway in *S. rosetta* will be necessary to uncover regulators upstream of Sr-p38 and if those regulators mirror function in animals or fungi.

Our approach of using sorafenib to uncover the role of *Sr*-p38 and other sorafenib-targeted kinases in *S. rosetta* allowed us to assess *Sr*-p38’s role before undertaking targeted genetics. Vertebrates express four p38 paralogs; the p38ɑ knockout is embryonic lethal, whereas knockouts for the other p38 paralogs are viable ^49^. *S. rosetta* is predicted to encode a family of stress-responsive kinases, including *Sr-*p38, *Sr-*JNK, and a homolog of ERK5 (EGD76774) (Figure S9A). We observed an increase in phosphorylation of a ∼45 kDa protein that was recognized by two independent phospho-p38 antibodies in *S. rosetta* cultures that were subjected to heat shock (Figures 4A-4B, S10A-S10B). The signal was lost in *Sr-p38*^1–15^ knockout strains (Figures 4B-4D, S12A-S12B), implicating *Sr-*p38, in the heat shock response. The viability of *Sr-p38*^1–15^ cells and the sensitivity of *Sr-*p38 mutants with reduced sorafenib binding (Figures 4F, S13C) suggest that sorafenib’s impact on cell proliferation is mediated by binding to other kinases.

Choanoflagellates have dynamic life histories and express diverse kinase families, including p38 kinases, that are found in animals ^8,10,17,43,62^. How kinases regulate additional aspects of choanoflagellate physiology, including life history transitions, remains to be investigated. We infer that fast-acting inhibition of kinase activity will be a powerful approach to study the roles of kinases that regulate cell state transitions in choanoflagellates. Because kinase inhibitors allow the enzymatic activity of kinases to be disrupted while preserving kinase localization and scaffolding functions, using small molecules as tools can distinguish whether kinase catalytic activity or other kinase functions are necessary for choanoflagellate development ^63,64^. Insight into the roles of individual kinases during the emergence of new cell-cell signaling networks (e.g. receptor tyrosine kinase signaling in the last common ancestor of animals, choanoflagellates, and their closest relatives ^8,10,13,62,65,66^) will be fundamental to understanding the contributions of kinase signaling to the origin of animal multicellularity.

## Materials and Methods

### Co-culturing of S. rosetta with the prey bacterium Echinicola pacifica

Choanoflagellates are bacterivores and require prey bacteria that are co-cultured in choanoflagellate media ^67^. *Echinicola pacifica*, a Bacteriodetes bacterium, grown in seawater-based media enriched with glycerol, yeast extract, and peptone sustains *S. rosetta* growth ^68^. This co-culture of *S. rosetta* and *E. pacifica*, publicly available from the American Type Culture Collection (ATCC) as ATCC PRA-390 and also known as “SrEpac” ^68^, was used in this study.

### High-throughput chemical screening in S. rosetta

To quantify changes in *S. rosetta* cell proliferation after small molecule treatment, we established a high-throughput screening pipeline.

We first assembled a library of 1255 compounds (Table S1) from commercial (Selleckchem Kinase Inhibitor Library Catalog #L1200 Lot: Z1316458) and academic (Kevan M. Shokat, University of California, San Francisco) sources. 98% of molecules in the library were characterized as human kinase inhibitors and 2% were compounds that are cytotoxic to other protists. Most of the kinase inhibitors in the library modulate human kinase activity by binding to the kinase active site and are ATP-competitive. Because we did not know if inhibitors designed to bind to human kinases would bind to choanoflagellate kinase homologs with the same potency or selectivity, we chose inhibitors with a range of selectivity: 75% of the human kinome is targeted by at least one inhibitor in the library, and the library includes inhibitors of all classified human kinase groups (Figure S1). Compounds dissolved at 10 mM in dimethyl sulfoxide (DMSO, Sigma #D8418) were placed into individual wells of 384-well deep well master plates (Corning #3342). We generated deep well stock plates with solutions that could be directly transferred to assay plates. Liquid handling (Agilent V11 Bravo) was used to dilute compound master plates (containing 10 mM compound in 100% DMSO) into deep well stock plates (containing 450 µM compound in 4.5% DMSO).

In a primary screen, *S. rosetta* cell counts were determined by analysis of acquired flow cytometry events after a 24-hour incubation. Assay plates were generated by plating 2 µL of the deep well stock plates into 384-well assay plates (Thermo Scientific #142761 using the Agilent V11 Bravo). 88 µL of SrEpac cultured in high-nutrient media (5% Sea Water Complete) ^69^ at exponential phase (∼9×10^5^ cells/mL) was diluted to 2×10^4^ cells/mL in high-nutrient media and dispensed into the assay plate (ThermoFisher Mulitdrop™ Combi with long standard dispensing tube cassette #24072677) to treat the SrEpac culture at 10 µM compound and 0.1% DMSO. After a 24-hour incubation, assay plates were individually placed into an autosampler (BD Biosciences High Throughput Sampler, HTS) running in high-throughput mode. 40 µL of each well was mixed twice (at 180 µL/second), and 10 µL of cells from each well were loaded onto a flow cytometer (BD Biosciences LSR II) at 1 µL/second. In between each well, the autosampler needle was washed with 400 µL of sheath fluid (1X phosphate-buffered-saline pH 7.4). Loaded cell samples were acquired on the cytometer with forward scatter (FSC) and side scatter (SSC) parameter voltages set to 538 and 308, respectively. A polygon gate from a DMSO well within a plate (D23 for Plate 1 and C23 for Plates 2-4 due to a shift in the distribution of observed events) was used to analyze events and quantify cell counts for all wells in an individual assay plate using FlowJo v10.8™ (BD Biosciences) (Figure S2A).

In a secondary screen, *S. rosetta* cell counts were determined by enumerating segmented cell objects from immunofluorescence images taken on an Opera Phenix high-content imager after a 48-hour incubation. Assay plates were generated by adding 4 µL of compound in the deep well stock plates into 96-well assay plates containing 176 µL of SrEpac in low-nutrient medium (1% Cereal Grass Medium and 1% Sea Water Complete ^70^) using an electronic multichannel pipetter (Rainin E4 Multi Pipette E8-20XLS+). Cells were initially expanded in high-nutrient media containing 4% Cereal Grass Medium and 4% Sea Water Complete ^70^ and at exponential phase (∼1×10^6^ cells/mL), diluted to 2×10^4^ cells/mL with AK-seawater. After a 48-hour incubation, cells were mixed in a thermomixer for two minutes at 800 rpm at room temperature to dislodge any cells attached to biofilm at the bottom of the 96-well plate. 100 µL of cells were transferred to Poly-D-Lysine (Sigma # P6407) coated 384-well imaging plates (Perkin Elmer Cell Carrier Ultra Plates #6057302) using an electronic multichannel pipetter (Rainin E4 Multi Pipette Multi E12-200XLS+). To optimize the screen, after 40 minutes of adherence, 1 µL of FM 1-43X mixture (1 µL of 500 µg/mL mixture of FM1-43X dye made by dissolving tube in 200 µL of methanol) was added to the cells and incubated for 15 minutes. 50 µL of the cell dye-mixture was removed followed by fixation and washing as described next for the full screen. For the full screen, after 40 minutes of adherence, 50 µL of cells were removed, and the remaining 50 µL were washed once with 50 µL 4X PBS. After incubating in the 4X PBS for 5 minutes, 50 µL was removed. Cells were fixed for 20 minutes at room temperature by adding 50 µL of 4% formaldehyde in PEM buffer (100 mM PIPES pH 7, 1 mM EGTA, 1 mM MgSO4). After fixation, cells were washed by removing 50 µL of solution from the plate and adding 50 µL of PEM buffer three times. After the final wash, 75 µL of solution was removed. At this point, cells for the optimization screen were imaged on a Perkin Elmer Opera Phenix with the following imaging specifications for the Fluorescein (FITC) channel: 20X water objective (NA 1.0, working distance 1.7mm, 646 µm^2^ field of view), 60ms and a three plane zstack at -8µm, -6µm and -4µm. For the full screen, after washing the fixative, cells were blocked by adding 75 µL of 2% BSA and 0.6% Triton-X100 in PEM for 30 minutes at room temperature. After blocking 25 µL was removed and 25 µL of primary antibody solution in 1% BSA and 0.3% Triton-X100 in PEM was added to stain the cell body and flagella overnight at 4°C (anti-tubulin, Abcam #ab6161, 1:1000 dilution). Due to the amount of time required for imaging, plates were staggered and processed one plate each day. On each imaging day, a plate was brought to room temperature and the primary antibody was washed three times by adding 50 µL of 1% BSA and 0.3% Triton-X100 in PEM and removing 50 µL of solution from the plate three times. Secondary antibody (Goat anti-rat Alexa Fluor 488, Invitrogen #A-11006, 1:300 dilution) and nuclear stain (DR<AQ5>, Thermo Scientific #62251, 1:500 dilution) were added in 25 µL of 1% BSA and 0.3% Triton-X100 in PEM and incubated for 2 hours at room temperature. After incubation, the secondary antibody was washed three times by adding 50 µL of PEM and removing 50 µL of solution from the plate three times. To stain the cell collar and obtain a second cell body marker, rhodamine phalloidin (Invitrogen #R415, 1:300 dilution) and Cell Tracker cmDIL (Invitrogen # C7000, 250 nM) were added for 25 minutes. To preserve the staining, 50 µL of solution was removed from the plate and 50 µL of 50% glycerol in PEM was added. 22 fields of view in 4 planes (each plane separated by 1 µm) of each well in the 384-well plate were imaged with a 40X Water immersion objective (NA 1.1, working distance 0.62mm, 323 µm^2^ field of view) on a Perkin Elmer Opera Phenix with optimized imaging specifications for each channel (Alexa 488 – Tubulin – 20ms; TRITC – Cell Tracker cmDIL / rhodamine phalloidin – 100ms; Brightfield – 100ms; Alexa Fluor 647 – DR<AQ5> / Nuclei – 1s). Due to restrictions on the available image area on the opera phenix, columns 1-23 of each 384-well plate were imaged first followed by a second scan with column 24 alone.

For both assays, the quantified cell counts were normalized to the average cell count of all DMSO wells within an individual plate to account for any plate-to-plate variation. Compounds were determined to significantly inhibit *S. rosetta* cell proliferation if the resulting normalized cell count had a p-value < 0.05 (based on two-tailed p-value calculated from z-score of all treated samples). The resulting normalized cell count data were plotted using GraphPad Prism 9.3.1™ (GraphPad San Diego, California, USA) (Figure 1; Figure S2B, Figure S2E, Figure S2G).

### Follow-up *S. rosetta* cell proliferation assays in response to compound treatment

Compound treatments post-library screening were conducted with commercially available inhibitors; sorafenib analogs previously synthesized ^40^; AD80 previously synthesized ^71^; and imatinib, PP1, and sunitinib provided by K. Shokat (UC San Francisco). Sorafenib (#S1040), regorafenib (#S5077), dasatinib (#S7782), PP2 (#S7008), and milciclib (#S2751) were purchased from Selleckchem. Glesatinib (#HY-19642A), masitinib (#HY-10209), lapatinib (#HY-50898), PP121 (#HY-10372), gliteritinib (#HY-12432), brigatinib (#HY-12857), RAF265 (#HY-10248), vemurafenib (#HY12057), skepinone-L (#HY-15300), BIRB 796 (#HY-10320), were purchased from MedChem Express. SU6656 (#13338) was purchased from Cayman Chemical. R406 (#A5880) was purchased from ApexBio Technlology.

SrEpac cultured in high-nutrient media (4% Sea Water Complete with 4% Cereal Grass) ^70^ in exponential phase (∼5×10^5^ - 9×10^5^ cells/mL) was diluted to lower density (1×10^4^ cells/mL), and 1 mL or 100 µL of cells were plated into 24-well or 96-well multiwell plates (Corning #3526, Thermo Scientific #260251), respectively. Cells were treated with compound by adding 1 µL of a 1000X compound stock in DMSO to a well in the 24-well plate or adding 1 µL of a 100X compound stock in DMSO. Equal volumes of DMSO were added to vehicle control wells so (v/v%) DMSO in controls was the same as compound treated wells. At set timepoints, cells were harvested. For cell assays in 24-well plates, cells in the well were pipetted up and down to resuspend the well and the well contents were transferred to a 1.5 mL Eppendorf tube. Cells were fixed by adding 40 µL of 37% formaldehyde (methanol-stabilized, Sigma-Aldrich #252549). For 96-well plates, 1 µL of 37% formaldehyde was added to each well by using a multichannel pipette, and a pierceable aluminum plate seal (USA Scientific TempPlate^®^ Sealing Foil) was added to cover the plate. The plate was vortexed at 2000rpm (24-well plates) or 3000rpm (96-well plates) in a plate vortexer (Eppendorf ThermoMixer C) to ensure equal fixation of cells in the well. Fixed cells were immediately counted or placed at 4°C for up to 2 weeks before counting. The cell density of sample timepoints along the full growth course was determined by analysis of micrographs taken on a Widefield microscope (Carl Zeiss AG Axio Observer.Z1/7, Oberkochen, Germany) ^70^, or brightfield imaging using a cell counter (Logos Biosystems LUNA-FL™). The resulting normalized cell density data at each timepoint was plotted using GraphPad Prism 9.3.1™ (GraphPad San Diego, California, USA) (Figure 2A; Figure 4F; Figure S6B-E; Figure S13B; Figure S14C). For comparisons between growth curves and phosphotyrosine signal (see *Assessment of* S. rosetta *kinase signaling by western blotting)* the <u>a</u>rea <u>u</u>nder the growth <u>c</u>urve (AUC) was analyzed using GraphPad Prism 9.3.1™ (GraphPad San Diego, California, USA) with baseline at Y=0 and minimum peak height > 10% above the baseline to maximum Y value (Figure S8; Figure S14B).

For dose-response assays, SrEpac cultured in high-nutrient media (5% Sea Water Complete ^69^ or 4% Sea Water Complete with 4% Cereal Grass ^70^) in exponential phase (∼5×10^5^ - 9×10^5^ cells/mL) was diluted to lower density. Starting density used varied based on treatment length: for 24-hour treatments and less, cells were plated at 2×10^5^ cells/mL; for 24-48 hour treatments, cells were plated at 1×10^5^ cells/mL; and for 48+ hour treatments, cells were plated at 5×10^4^ cells/mL. Cell density was determined at the treatment endpoint using the same approach as the treatment growth curves described in the preceding paragraph. The resulting normalized cell density data at each dose was plotted using GraphPad Prism 9.3.1™ (GraphPad San Diego, California, USA). (Figure 2B; Figure S2C; Figure 13C; Figure S14D)

### Live imaging of treated S. rosetta cultures

Cells were imaged by differential interference contrast (DIC) using a 100× (oil immersion, Plan-Apochromat, 1.4 NA) Zeiss objective mounted on a Zeiss Observer Z.1 with a Hamamatsu Orca Flash 4.0 V2 CMOS camera (C11440-22CU). Movies were annotated using the Annotate_movie^72^ plugin on Fiji (v 2.3.0/1.53q)^73^ (Movies S1-S4).

### Assessment of *S. rosetta* kinase signaling by western blotting

After compound treatment, the SrEpac culture was harvested and lysed to quantify protein abundance and immunoblotting by western blot. *S. rosetta* cells were harvested by centrifugation at 6,000g for five minutes in Falcon tubes in a swinging bucket centrifuge and transferred into 1.5 mL Eppendorf tubes and washed two times with 4X phosphate buffered saline (PBS, 6.2 mM potassium phosphate monobasic, 621 mM sodium chloride, 10.8 mM sodium phosphate dibasic) with centrifugation at 6,000g for 5 minutes in a fixed angle centrifuge at room temperature in between each wash. Cells were resuspended and lysed in digitonin lysis buffer (20 mM Tris pH 8, 150 mM potassium chloride, 5 mM magnesium chloride, 250 mM sucrose, 1 mM Pefablock® SC serine protease inhibitor (Sigma-Aldrich Cat# 76307), 8 mM digitonin, 1 mM dithiothreitol, 0.06 U/µL benzonase nuclease, 1X Roche PhosSTOP phosphatase inhibitor cocktail, 1X Roche cOmplete protease inhibitor cocktail) for 30 minutes. Lysed cells were spun at 18,000g for 15 minutes at 4°C and supernatants were isolated. Protein concentration in supernatants was determined by Bradford assay. Samples were boiled in loading dye (LiCOR #928-40004) and equal protein amounts for each sample were loaded onto NuPAGE™ 4-12% Bis-Tris SDS-PAGE gels (Invitrogen Cat#s WG1402BOX, WG1403BOX, NP0335BOX, NP0336BOX). To resolve bands recognized by the phospho-p38 antibody (Promega #V1211), a NuPAGE™12% Bis-Tris SDS-PAGE gel (Invitrogen Fisher NP0349BOX) was used. PageRuler™ or PageRuler Plus™ prestained protein ladder (Thermo Scientific #26614 and #26620), EGF-stimulated A431 cell lysate control (Sigma Aldrich #12-302), and *E pacifica* control lysate (lysed as for *S. rosetta* cells) were added to wells and gels were run in Novex NuPAGE MES buffer (Invitrogen Cat# NP000202). Gels were transferred to 0.45 µm nitrocellulose (Bio-Rad) in Tris-Glycine Buffer (Bio-Rad #1610734) with 10% Methanol. After transfer, blots were stained with LI-COR Revert™ 700 and total protein imaged. After total protein was stained, blots were blocked with 5% bovine serum albumin in Tris buffered saline (TBS, 25 mM Tris, 150 mM sodium chloride) for 1 hour. Primary antibodies (see next paragraph) were added in TBS with Tween (0.1%) and left overnight. Blots were washed with TBS-Tween four times. LI-COR secondary IRDyes® were added in TBS-Tween at 1:10,000 dilution and incubated for 1 hour at room temperature. Blots were washed with TBS-Tween four times followed by TBS one time and imaged with LI-COR Odyssey® imager (Figure 2C; Figure 4D, 4E, 4G; Figure S2D; Figure S6A; Figure S7; Figure S10; Figure S12; Figure S14E, Figure S15). Staining intensities were quantified with Image Studio Lite 5.2.5 (LI-COR, 2014) (Figure 4B, 4C; Figure S2D; Figure S10A; Figure S12A, S12B; Figure S14E) and the resulting normalized cell density data at each dose was plotted using GraphPad Prism 9.3.1 μ (Figure 4B, 4C).

Primary antibodies for pY1000 (#8954), anti-phospho-Erk (anti-phospho p44/42 MAPK #4370), anti-phospho(Ser/Thr)-Phe (#9631), anti-phosphotheronine (#9381) and anti-phosphothreonine-proline (#9391) were purchased from Cell Signaling Technology. Anti-phospho p38 MAPK (Anti-ACTIVE® p38, #V1211) was purchased from Promega or Biorad (p38 MAPK pThr180/pTyr182, #AHP905). Anti-phosphoserine (Rb X, #AB1603) was purchased from Millipore Sigma. Anti-alpha-tubulin (YOL 1/34, #ab6161) was purchased from Abcam.

### ActivX Mass Spectrometry workflow to identify *S. rosetta* proteins that bind sorafenib

For all mass spectrometry experiments, SrEpac cultures were grown to a high density in Pyrex baffled flasks without shaking. In tall necked flasks, cultures were grown in the maximum volume of culture media, and an aquarium pump was used to bubble air into the foam-plugged PYREX® Delong flask pierced with a serological pipette at a bubbling rate of approximately one bubble/second ^74^. For wide 2.8L PYREX® Fernbach flasks, bubbling was not needed. *S. rosetta* cells were harvested by spinning in 200mL Nalgene conicals at 2000g for 10 minutes in a swinging bucket centrifuge at room temperature to obtain cell pellets. Cells were first washed by resuspending cell pellets from four conicals in approximately 45 mL of 4X PBS, transferring cells to 50 mL Falcon tubes, and spinning at 2000g for 10 minutes in a swinging bucket centrifuge at room temperature. Cells were washed a second time by resuspending cells in ∼15 mL of 4X PBS per 50 mL Falcon tube, transferring cells to 15 mL Falcon tubes, and spinning at 2000g for 10 minutes in a swinging bucket centrifuge at room temperature. Cells were then ready for the ActivX probe workflow.

For ActivX probe enrichment, cells were lysed and 500 µL of 5 mg/mL supernatants were obtained as previously described ^42^. 20 mM of manganese chloride cofactor was added to the lysate and incubated for 5 minutes followed by the addition of 100 µM sorafenib or 1% DMSO (vehicle control) and incubation for 10 minutes. 20 µM ActivX probe was added for kinase capture. Biotinylated proteins were captured using streptavidin beads in the presence of 6M urea / immunoprecipitation (IP) lysis buffer, and samples were washed with 6M urea / IP lysis buffer. Protein samples were provided to the University of California, Davis mass spectrometry facility (https://cmsf.ucdavis.edu) on beads and underwent standard tryptic digestion with Promega ProteaseMAX™ Surfactant, Trypsin Enhancer. Briefly, samples were first reduced at 56°C for 45 minutes in 5.5 mM DTT followed by alkylation for one hour in the dark with iodoacetamide added to a final concentration of 10 mM. Trypsin was added at a final enzyme:substrate mass ratio of 1:50 and digestion carried out overnight at 37°C. The reaction was quenched by flash freezing in liquid nitrogen and the digest was lyophilized. Digest was reconstituted in 0.1% TFA with 10% acetonitrile prior to injection. For quantification of peptides within each sample, 500 fmol of Hi3 *E. coli* standard reagent ClpB (Waters, Milford, MA) was placed into each sample and injected with 1.0 ug of total digest. Each sample was run in triplicate.

The mass spectrometry instrument used to analyze the samples was a Xevo G2 QTof coupled to a nanoAcquity UPLC system (Waters, Milford, MA). Samples were loaded onto a C18 Waters Trizaic nanotile of 85 µm × 100 mm; 1.7 μm (Waters, Milford, MA). The column temperature was set to 45°C with a flow rate of 0.45 mL/min. The mobile phase consisted of A (water containing 0.1% formic acid) and B (acetonitrile containing 0.1% formic acid). A linear gradient elution program was used: 0–40 min, 3–40 % (B); 40-42 min, 40–85 % (B); 42-46 min, 85 % (B); 46-48 min, 85-3 % (B); 48-60 min, 3% (B). Mass spectrometry data were recorded for 60 minutes for each run and controlled by MassLynx 4.2 SCN990 (Waters, Milford, MA). Acquisition mode was set to positive polarity under resolution mode. Mass range was set form 50 – 2000 Da. Capillary voltage was 3.5 kV, sampling cone at 25 V, and extraction cone at 2.5 V. Source temperature was held at 110C. Cone gas was set to 25 L/h, nano flow gas at 0.10 Bar, and desolvation gas at 1200 L/h. Leucine–enkephalin at 720 pmol/µl (Waters, Milford, MA) was used as the lock mass ion at *m*/*z* 556.2771 and introduced at 1 µL/min at 45 second intervals with a 3 scan average and mass window of +/-0.5 Da. The MSe data were acquired using two scan functions corresponding to low energy for function 1 and high energy for function 2. Function 1 had collision energy at 6 V and function 2 had a collision energy ramp of 18 − 42 V.

RAW MSe files were processed using Protein Lynx Global Server (PLGS) version 3.0.3 (Waters, Milford, MA). Processing parameters consisted of a low energy threshold set at 200.0 counts, an elevated energy threshold set at 25.0 counts, and an intensity threshold set at 1500 counts. Each sample was searched against the *Salpingoeca rosetta* genome hosted on Ensembl Genomes ^17,75^. Each databank was randomized within PLGS and included the protein sequence for ClpB. Possible structure modifications included for consideration were methionine oxidation, asparagine deamidation, glutamine deamidation, serine dehydration, threonine dehydration, and carbamidomethylation of cysteine. For viewing, PLGS search results were exported in Scaffold v4.4.6 (Proteome Software Inc., Portland, OR). We quantified absolute protein abundance ^76^ and focused our attention on kinases that were present in both DMSO and sorafenib pretreated samples but were enriched in multiple DMSO replicates, represented by higher PLGS scores ^76^. The resulting data was plotted with GraphPad Prism 9.3.1 μ (Figure 3).

### Generation of Sr-p38^1–15^ and Sr-JNK^1–15^ strains by genome editing

Candidate guide RNA sequences that targeted early in the *Sr-p38* and *Sr-JNK* open reading frame were identified using the EuPaGDT tool (http://grna.ctegd.uga.edu/) and the *S. rosetta* genome ^17^ hosted on Ensembl Protists (Ensembl 108) ^77^. Guide RNA length was set at 15 and an YRNGRSGGH PAM sequence was used. Guide RNA candidates were filtered for guides with one on-target hit (including making sure the guides do not span exon-exon boundaries), zero off-target hits (including against the genome of the co-cultured bacterium *E. pacifica*), lowest strength of the predicted secondary structure (assessed using the RNAfold web server: http://rna.tbi.univie.ac.at/cgi-bin/RNAWebSuite/RNAfold.cgi), and annealing near codon 15 of *Sr-p38* and *Sr-JNK*. A crRNA with the guide sequence TGCAAGTCTGTGTAGCACGA for *Sr-p38* and TTGTCGATGTGTGGAAGCAG for *Sr-JNK*, as well as universal tracrRNAs, were ordered from IDT (Integrated DNA Technologies, Coralville, IA). Repair templates were produced by PCR using a previously published plasmid encoding the pEFL-pac-5’Act (Addgene ID pMS18, catalog number #225681) cassette as a template. The premature termination sequence (5’-TTTATTTAATTAAATAAA-3’) was included in custom primers with 50 bp homology arms to the *Sr-p38* or *Sr-JNK* locus. PCR reactions were set up in 50 μL reactions using Q5 high fidelity DNA polymerase with 50 ng plasmid as template, 200 μM dNTPs, 0.25 μM of each primer and 0.02 U/ μL Q5 polymerase. The following PCR program was run on an Applied Biosystems Veriti 96-well Thermal Cycler: 30” 98°C; 40X (10” 98°C; 30” 68°C; 1’ 72°C); 2’ 72°C. The PCR product size (expected 2 kb) was visually checked by running 2 μL of the reaction on a 1% (w/v) agarose gel containing ethidium bromide at 1 µg/mL run in TAE buffer and visualized with the Alpha Innotech 2000 Photo Imaging System Ultraviolet Transilluminator (Figure S11A, S11B). PCR products were purified using a PCR purification kit (NEB Monarch Cat #T1130L and SydLabs Tiniprep columns Cat# MB0110L) and the final product was eluted with pre-warmed 20 μL milliQ water which was left to incubate on the column for 10 minutes before eluting. 1.5 μL was used to measure DNA concentration using a NanoDrop spectrophotometer (ThermoFisherScientific NanoDrop 2000). 6 μg of the remaining DNA was concentrated in 2 μL by evaporation for 2 hours at 55°C or lyophilized and resuspended into 2 μL warm milliQ water and then used as a repair template for nucleofection. Genome editing proceeded as described previously ^57^, but with 3H Buffer (75 mM Tris-HCl, 20 mM HEPES, 90 mM NaCl, 15 mM MgCl2, 5 mM KCl, 10 mM glucose, 0.4 mM calcium nitrate) replacing Lonza SF buffer. Puromycin (80 μg/mL) was added after 24 hours. At 72 hours post addition of puromycin, the wild-type cells had died, and puromycin-resistant cells appeared for *Sr-JNK*^1–15^ nucleofections. Puromycin-resistant cells for *Sr-p38*^1–15^ appeared at 96 hours. Genotyping primers used to confirm insertion of the premature termination sequence and puromycin resistance cassettes^57^ were AACAGGAGGCACAGTTACGA (forward) and GAACAAGCAACACACCACCA (reverse) for *Sr-p38*; CGTTAATCGACGACGCCAA (forward) and ATGAGCTGGATGTGGGGGA (reverse) for *Sr-JNK*. Premium PCR Sequencing was performed by Plasmidsaurus using Oxford Nanopore Technology with custom analysis and annotation to genotype clones. Base calls in the insertion region for *Sr-p38*^1–15^ are in Figure S11C and *Sr-JNK*^1–15^ in S11D.

### Generation of Sr-p38^T110M^ strains by genome editing

Candidate guide RNA sequences were obtained for *Sr-p38* using the EuPaGDT tool (http://grna.ctegd.uga.edu/) and the *S. rosetta* genome ^17^ hosted on Ensembl Protists (Ensembl 108) ^77^ as previously published ^70^. Guide RNA length was set at 15, and an NGG PAM sequence was used. Guide RNA candidates were filtered for guides with one on-target hit (including making sure the guides do not span exon-exon boundaries), zero off-target hits (including against the genome of the co-cultured bacterium *E. pacifica*), lowest strength of the predicted secondary structure (assessed using the RNAfold web server: http://rna.tbi.univie.ac.at/cgi-bin/RNAWebSuite/RNAfold.cgi), and annealing near codon 110 of *Sr-p38*. A crRNA with the guide sequence CTACATCATCACAGAGAAGA, as well as universal tracrRNAs, were ordered from IDT (Integrated DNA Technologies, Coralville, IA). Repair templates were designed as single-stranded DNA oligos, in the same sense strand as the guide RNA, with 50 base pairs of genomic sequence on either side of the DSB cut site. The repair oligo GGTTGACCTGTACATCTCCGACGCGCGTGACATCTACATCATCATGGAGAAGATGGTGTGTATCTTTGACGCGGGTTGACTGGCCGTGATGGCGCGTGTT was ordered from IDT as an Ultramer. Genotyping primers were AGATTCGTTCCAGCGGAATACA (forward) and GGGAAGAAGTGCGGAGTGAA (reverse). Genome editing proceeded as described previously ^70^. Clones were Sanger sequenced at the UC Berkeley DNA Sequencing Facility.

### Bioinformatic analysis of the S. rosetta kinome

The *S. rosetta* kinome was annotated based on previously predicted kinases ^17^ and orthoDB ortholog annotation ^78^ of *S. rosetta* and human protein sequences in Uniprot ^79^ (Figure S1). For prediction of choanoflagellate p38 kinases, a HMMER profile was generated from human and previously predicted *S. rosetta* p38 kinases ^17,43^ and searched against available genomes and transcriptomes of choanoflagellates ^9,12,80,81^ (Figure S6A-B). Protein targets of individual human kinase inhibitors were manually annotated and plotted using CORAL ^82^ (Figure S1). References for kinase inhibitory data for each compound is available in Table S1. To analyze conservation within the kinase domain, kinase sequence alignments of predicted kinases were generated with Clustal Omega ^83^ (Figure S5B-C; Figure S9B) and amino acid logos were generated with WebLogo ^84^ (Figure S5A).

## Supporting information

Movie S1 - Glesatinib

Movie S2 - Sorafenib

Movie S3 - Sorafenib

Movie S4 - DMSO

## Author Contributions

F.U.R., M.C., A.C.D., and N.K. designed research; F.U.R. performed screening and choanoflagellate research, F.U.R. M.C., M.H.T.N., and I.H. generated choanoflagellate strains, and A.P.S. synthesized sorafenib analogs; F.U.R., A.C.D., and N.K. analyzed data; and F.U.R., and N.K. wrote the paper.

## Conflicts of Interest

There are no conflicts of interest to declare.

## Data Availability

Data supporting this article are included in the Supplementary Information. Raw data for this article, including flow cytometry fcs files, genotyping, image segmentation output files, sequence alignments, raw counts and cell densities, mass spectrometry peptides, raw western blot images, western blot cropping and western blot quantitation, are available at Figshare at https://doi.org/10.6084/m9.figshare.20669730.v3.

## Acknowledgments

We thank D. Booth, M. Carver, A. Garcia De Las Bayonas and J. Reyes-Rivera for critical reading of the manuscript; K. Shokat (University of California, San Francisco) for the contribution of kinase inhibitors in the screened library and Shokat lab members for discussions; M. West and P. He of the High-Throughput Screening Facility (QB3 HTSF) at UC Berkeley who provided access to the Agilent V11 Bravo liquid handling system for screening plate generation through funding from the National Institutes of Health (S10OD021828); H. Nolla, A. Valeros and the CRL-FACS facility and staff at UC Berkeley for their support and providing access to the BD LSRII flow cytometer and BD HTS sample loading robot; A. Andaya of the mass spectrometry facility at UC Davis. The A.C.D. laboratory receives support from the NIH (RO1s CA227636, CA258736, CA256480, and R56 AG066712), the Mark Foundation for Cancer Research (20-030-ASP, 21-039-ASP) and Alex’s Lemonade Stand Foundation for Childhood Cancer. N.K. and F.U.N.R.’s work was supported by an Investigator award (N.K.) and the Hanna H. Gray Fellows Program (F.U.N.R.), both from the Howard Hughes Medical Institute.

## Notes

https://doi.org/10.6084/m9.figshare.20669730.v3

### Competing Interest Statement

The authors have declared no competing interest.

### Summary of Updates

This version of the manuscript has been revised to include additional experiments that were conducted as a part of the study. These experiments appear in the main text and supplementary information. The author list was updated to include new authors who conducted portions of these experiments.

