## Supplementary Information for "A stress-responsive p38 signaling axis in choanoflagellates"

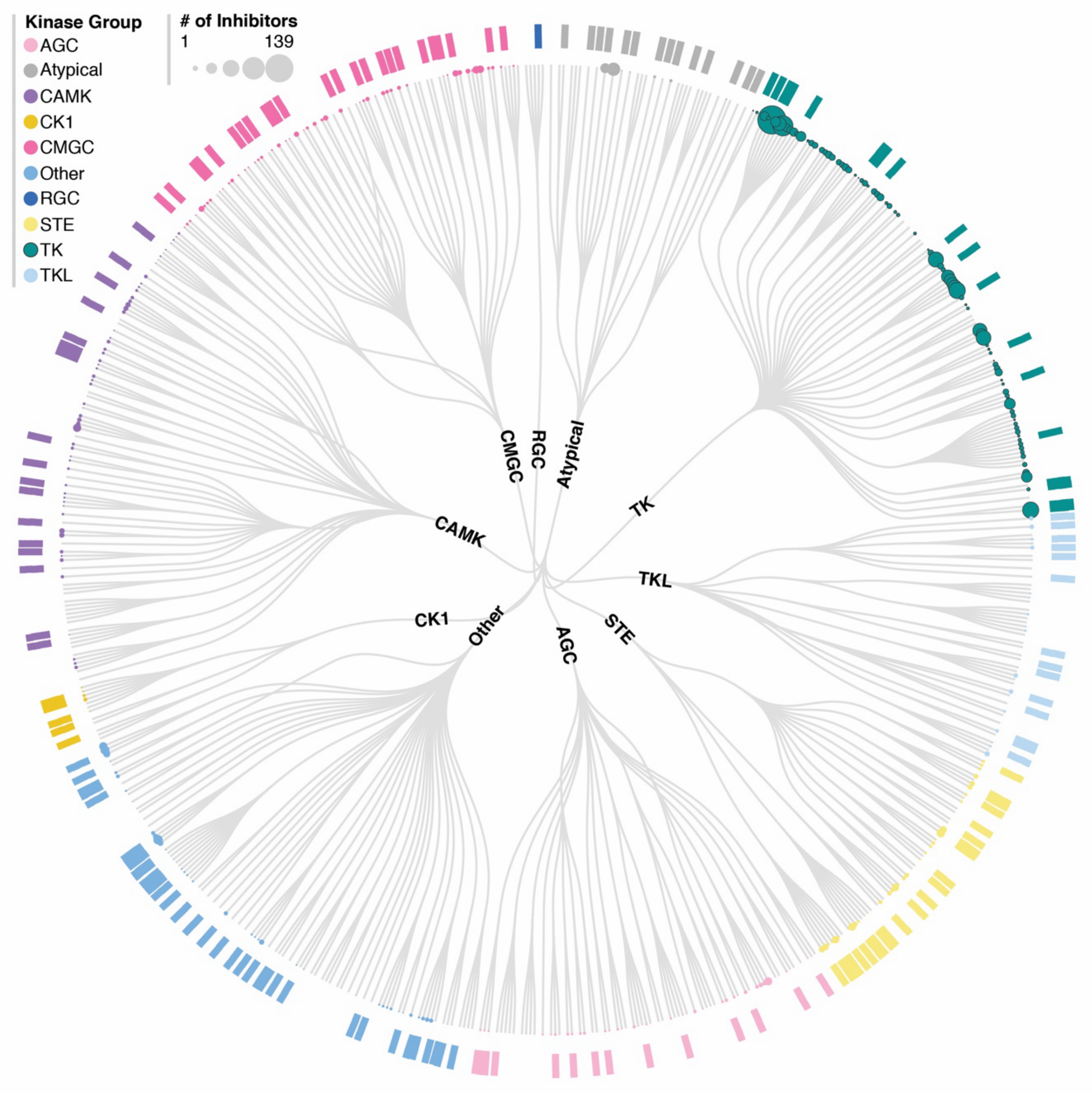


**Figure S1. The screened kinase inhibitor library includes targets of all human kinase groups and is predicted to have broad coverage for the previously annotated *S. rosetta* kinome.** The screening library consists of small molecule kinase inhibitors with broad coverage of the human kinome. Protein kinase targets of inhibitors in the library are indicated as colored outer nodes in the radial dendrogram. Node color indicates kinase group and node size indicates the number of inhibitors for a kinase target. The predicted presence of a *Salpingoeca rosetta* kinase homolog [1] is indicated with an outermost bar. Radial dendrogram made with CORAL [2].

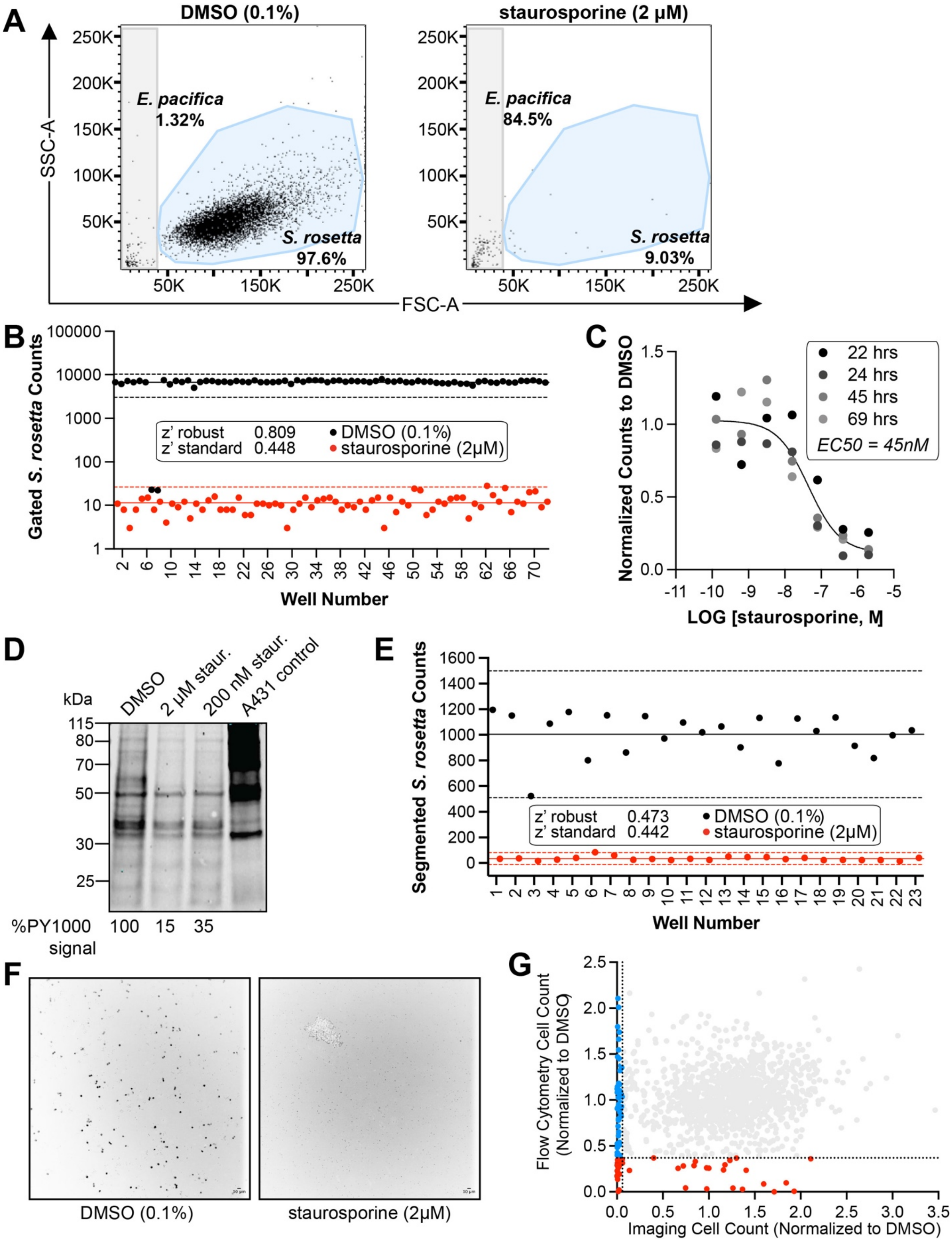
A
DMSO (0.1%)
staurosporine (2 μM)
SSC-A
FSC-A
250K
200K
150K
100K
50K
E. pacifica
1.32%
S. rosetta
97.6%
E. pacifica
84.5%
S. rosetta
9.03%
B
Gated S. rosetta Counts
100000
10000
1000
100
10
1
z' robust 0.809
z' standard 0.448
DMSO (0.1%)
staurosporine (2μM)
Well Number
C
Normalized Counts to DMSO
22 hrs
24 hrs
45 hrs
69 hrs
EC50 = 45nM
LOG [staurosporine, M]
D
kDa
DMSO
2 μM staur.
200 nM staur.
A431 control
115
80
70
50
30
25
%PY1000 signal
100
15
35
E
Segmented S. rosetta Counts
z' robust 0.473
z' standard 0.442
DMSO (0.1%)
staurosporine (2μM)
Well Number
F
DMSO (0.1%)
staurosporine (2μM)
G
Flow Cytometry Cell Count (Normalized to DMSO)
Imaging Cell Count (Normalized to DMSO)

**Figure S2. Staurosporine shows dose-responsive inhibition of *S. rosetta* tyrosine phosphorylation and cell proliferation that is quantifiable by high-throughput flow cytometry and imaging.**

(A) *S. rosetta* cells can be distinguished from their bacterial prey, *Echinicola pacifica* (Supplementary Materials and Methods) by forward scatter area (FSC-A) vs. side scatter area (SSC-A) analysis. *E. pacifica* and *S. rosetta* populations were gated after a 48-hour treatment with vehicle (DMSO, 0.1%) or staurosporine (2 μM) in a 384-well plate. The grey rectangle designates the *E. pacifica* gate and the blue polygon designates the *S. rosetta* gate used to compare treated multiwells. The percentage indicated in gate label is the percentage of events within each well that correspond to each gate. In vehicle (DMSO) wells, most events fell within the *S. rosetta* cell gate and in staurosporine treated wells, a majority of events were analyzed as *E. pacifica* cells, indicating a decrease in *S. rosetta* cell population with staurosporine treatment.

(B) The clear separation between cell counts quantified from vehicle (DMSO, 0.1%) and staurosporine (2 μM) treated wells demonstrated that flow cytometry analysis of treated *S. rosetta* cultures can accurately identify compounds, screened with one replicate, that inhibited *S. rosetta* cell proliferation. The confidence in being able to distinguish vehicle from staurosporine treated wells was quantified by z', which shows acceptable assay statistics for biological screens (z' standard close to 0.5 and z' robust > 0.5) [3,4]. The solid line indicates the treatment mean, and dashed lines indicate three standard deviations above and below the treatment mean with line colors corresponding to DMSO vehicle (black) and staurosporine treated wells (red). The red dashed line three standard deviations below the staurosporine treated wells is not present on the graph because the value was below one and the y-axis is plotted on a $\log_{10}$ scale.

(C) Staurosporine treatment was dose-responsive, and inhibition of *S. rosetta* cell proliferation was observable within 22 hours. Dose-response data are from single treatments of *S. rosetta* cultures with staurosporine at four different endpoints. One nonlinear regression curve fits all replicates as determined by an extra sum-of-squares F Test ($p < 0.001$).

(D) Staurosporine treatment inhibited *S. rosetta* kinase activity. Western blot analysis of *S. rosetta* cultures treated with staurosporine for 10 min. at 2 μM and 200 nM showed a reduction in phosphotyrosine (pY1000) signal relative to the DMSO (vehicle) control. Lysate from EGF-stimulated mammalian A431 cell line was used as a positive control for phosphotyrosine (pY1000). Total protein stain was used to normalize protein loaded for quantification of % pY1000 signal.

(E) Cell counts determined by segmentation of images of stained *S. rosetta* cells demonstrated that imaging could also distinguish vehicle from staurosporine treated wells. The confidence in being able to separate staurosporine treated wells from vehicle was quantified by z', which shows assay statistics worse than flow cytometry but still separated by three standard deviations above and below the treatment means. Line colors corresponding to DMSO vehicle are indicated in black and staurosporine treated wells are indicated in red.

(F) Representative images from one out of nine frames of a vehicle (DMSO, 0.1%) and a staurosporine (2 μM) treated well. Each image is a maximum-intensity projection of three planes of cells stained with FM 1-43FX. The lookup table (LUT) of each image was inverted for ease of visualization. DMSO treated wells show round cells that are easily quantified whereas staurosporine treated wells only show cell debris. Scale = 10 μm.

(G) Comparison of flow cytometry and imaging cell counts (Table S1). Compounds identified by flow cytometry are indicated in red and compounds identified by imaging are blue. Significance was determined based on two-tailed p-value < 0.05 calculated from z-score). Compounds that showed no effect are light grey.

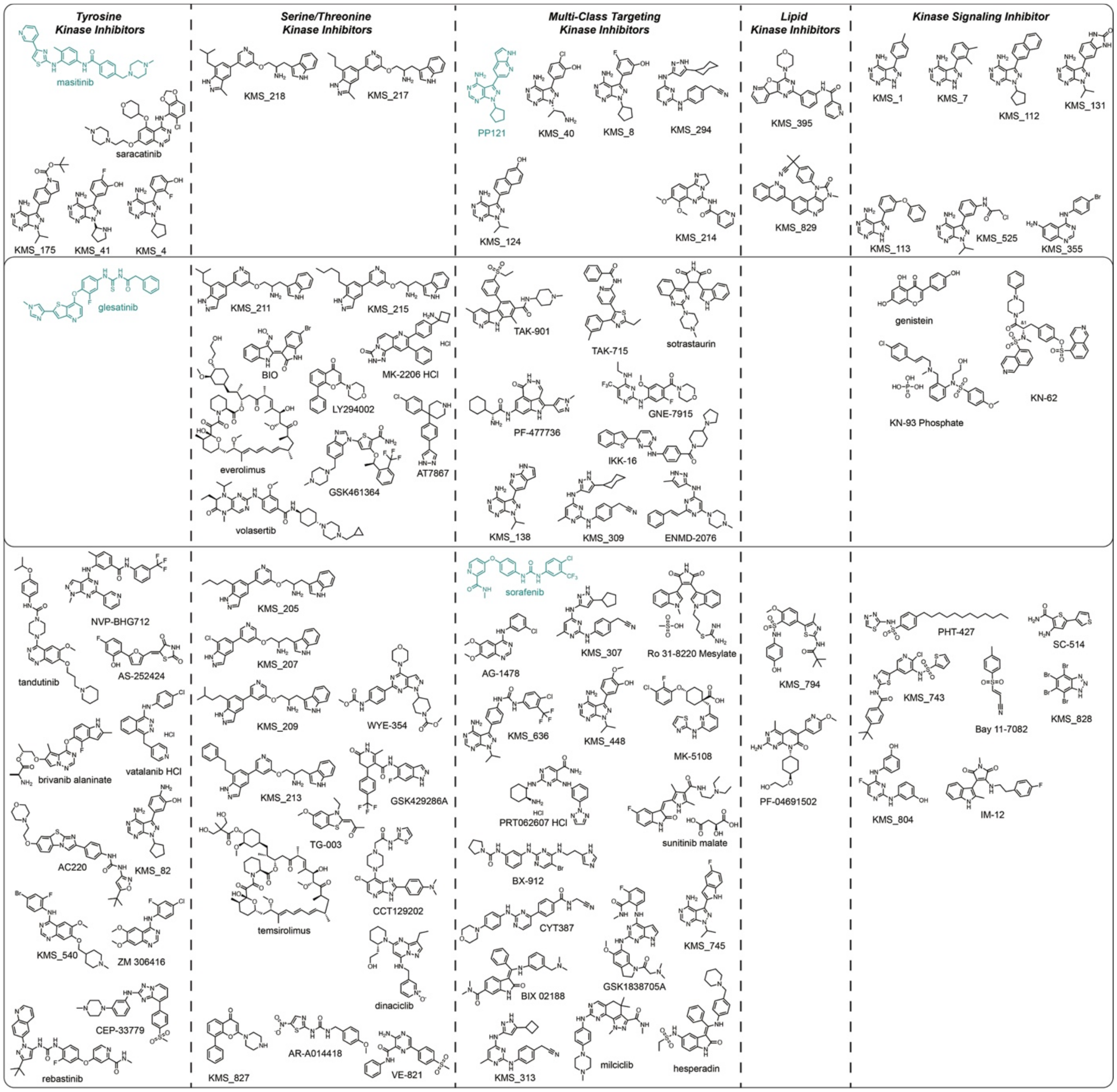


**Figure S3. Chemical structures of inhibitors of *S. rosetta* cell proliferation.** Inhibitors (Figure 1, red dots – flow cytometry, blue dots – imaging) are grouped by specificity towards kinase class (biochemical inhibition of human kinase within group with $IC_{50}$ < 100 nM) or kinase pathway (inhibit kinase signaling but do not directly bind to a kinase or have an unannotated kinase target). Four molecules (masitinib, PP121, glesatinib, sorafenib) that inhibit human tyrosine kinases and were further studied to assess impacts on *S. rosetta* phosphosignaling are indicated in teal.

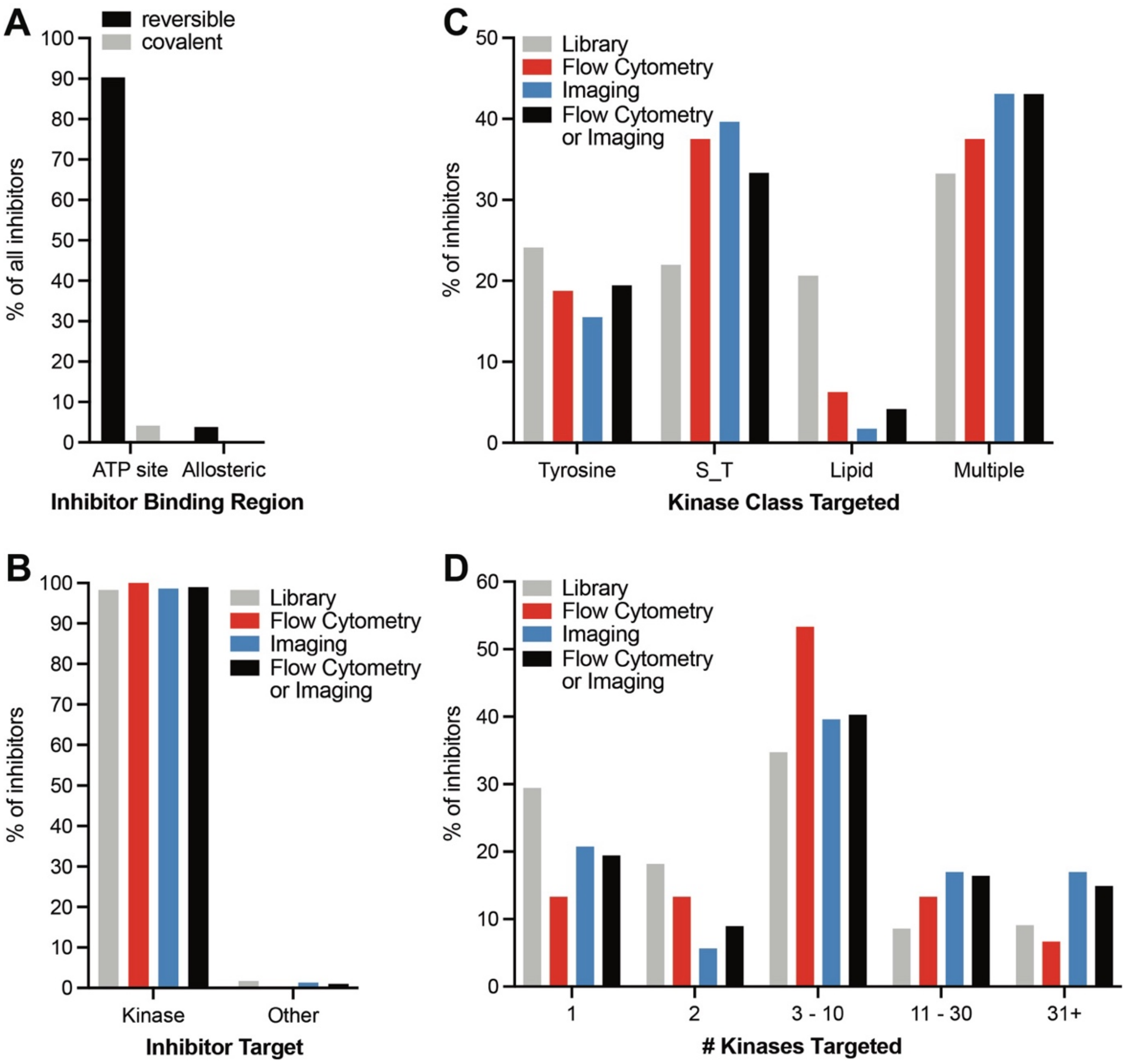


**Figure S4. Characterized mechanism of action, binding modes, and human kinase selectivity features of the 95 inhibitors of *S. rosetta* cell proliferation identified by flow cytometry.**

(A) Most kinase inhibitors in the library are reversible and ATP-competitive.

(B) Most compounds in the library and the identified inhibitors of *S. rosetta* cell proliferation are kinase inhibitors.

(C) Inhibitors of *S. rosetta* proliferation have potent biochemical inhibition ($IC_{50} < 100$ nM) of human kinases from all human kinase classes. Inhibitors of *S. rosetta* cell proliferation were modestly enriched for activity against human tyrosine (TK) and serine-threonine (S_T) kinases.

(D) Compounds that inhibit *S. rosetta* proliferation range from selective to non-selective for inhibition of human kinases. Inhibitors of *S.* rosetta proliferation were enriched for molecules with more than two human kinase targets.

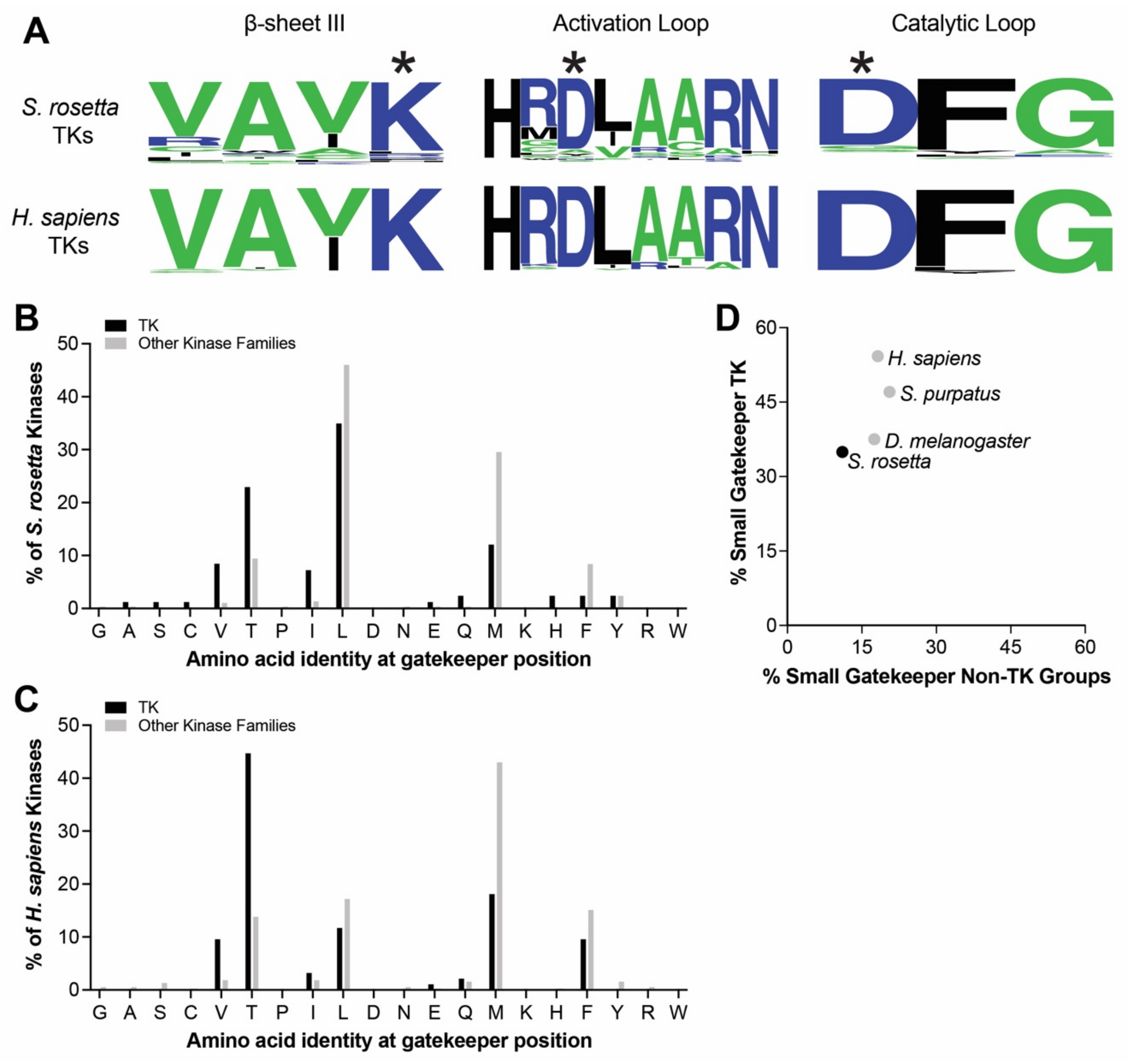


**Figure S5. Active site residues necessary for kinase activity and tyrosine kinase inhibitor selectivity (gatekeeper) are conserved in *H. sapiens* and *S. rosetta* kinases.**

(A) Sequence logos indicate conserved amino acids in *S. rosetta* and *H. sapiens* tyrosine kinase active sites. Amino acids are colored based on being charged at neutral pH (blue), having small size (green) or being any other amino acid (black). Amino acids indicated by (*) are required for enzymatic activity in studied human protein kinases [5,6].

(B) *S. rosetta* tyrosine kinase gatekeepers (black) are enriched for small amino acids compared to other *S. rosetta* kinase families (grey). Gatekeepers of ten *S. rosetta* TKs could not be assigned due to incomplete sequence.

(C) *H. sapiens* tyrosine kinase gatekeepers (black) are enriched for small amino acids compared to other kinase families (grey). Pseudokinase domains of *H. sapiens* JAK family kinases are not included.

(D) Small gatekeeper amino acids (G, A, S, C, V, T) are enriched in tyrosine kinases compared to other kinase groups in animals (represented by *H. sapiens, D. melanogaster, C. elegans, S. purpatus, A. queenslandica*) and *S. rosetta*.

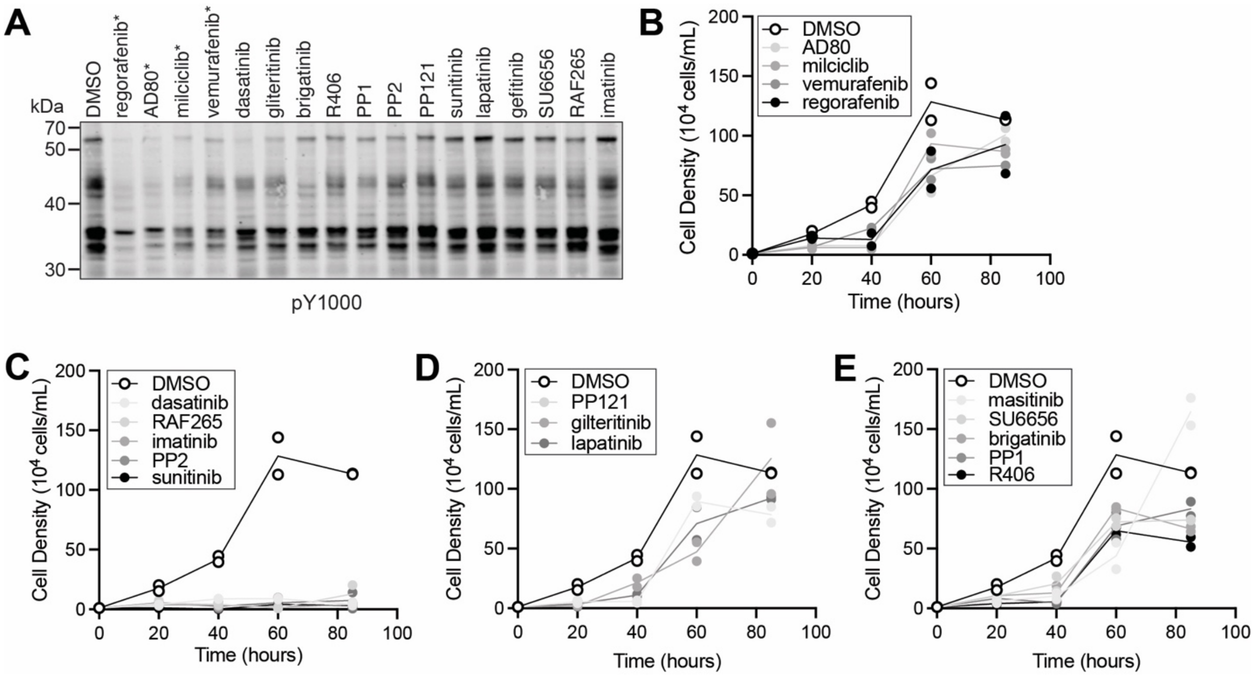


**Figure S6. Inhibition of *S. rosetta* phosphotyrosine signaling is not always connected to inhibition of *S. rosetta* cell proliferation.**

(A) *S. rosetta* cultures were treated with a panel of small molecules that inhibit human tyrosine kinases at 1 µM for 15 minutes and assessed for changes in tyrosine phosphorylation. The treated lysates were analyzed by western blot with a phosphotyrosine antibody (pY1000) to identify if any changes in tyrosine phosphorylation occurred. Treatment of *S. rosetta* cells with compounds indicated by "*" generated lysates that showed a global decrease in phosphotyrosine staining. The resulting level of tyrosine phosphorylation of proteins at ~60kDa, ~45kDa and ~35kDa is indicated by arrows. No changes in relative serine or threonine phosphorylation were observed by western blot (Fig. S7).

(B) *S. rosetta* cultures treated with 1 µM AD80, milciclib, vemurafenib, and regorafenib had reduced cell density at 40 hours in comparison to vehicle (DMSO) control but were not significant (p-value <0.05) after 40 hours. Two biological replicates were conducted per treatment, and each point represents the mean of three measurements from each biological replicate. For timepoints at 40, 60, and 85 hours, cell densities of inhibitor-treated cultures were considered significant if different from vehicle (DMSO) (p-value <0.05). Significance was determined by a two-way ANOVA multiple comparisons test.

(C) *S. rosetta* cultures treated with 1 µM imatinib and PP2 had reduced cell density at 40 hours in comparison to vehicle (DMSO) control but PP2 was not significant (p-value <0.05) after 40 hours. *S. rosetta* cultures treated with 1 µM dasatinib and sunitinib had reduced cell density at 85 hours in comparison to vehicle (DMSO) control. Treatment with RAF265 did not reach significance (p=0.0666 at 40 hours and 0.0847 at 85 hours. Two biological replicates were conducted per treatment, and each point represents the mean of three measurements from each biological replicate. For timepoints at 40, 60, and 85 hours, cell densities of inhibitor-treated cultures were considered significant if different from vehicle (DMSO) (p-value <0.05). Significance was determined by a two-way ANOVA multiple comparisons test.

(D) *S. rosetta* cultures treated with 1 µM PP121, gilteritinib, and lapatinib did not show reduced cell density at in comparison to vehicle (DMSO) at all timepoints tested. Two biological replicates were conducted per treatment, and each point represents the mean of three measurements from each biological replicate. For timepoints at 40, 60, and 85 hours, cell densities of inhibitor-treated cultures were considered significant if different from vehicle (DMSO) (p-value <0.05). Significance was determined by a two-way ANOVA multiple comparisons test.

(E) *S. rosetta* cultures treated with 1 µM masitinib, SU6656, brigatinib, PP1, and R406 did not show reduced cell density at in comparison to vehicle (DMSO) at all timepoints tested. Two biological replicates were conducted per treatment, and each point represents the mean of three measurements from each biological replicate. For timepoints at 40, 60, and 85 hours, cell densities of inhibitor-treated cultures were considered significant if different from vehicle (DMSO) (p-value <0.05). Significance was determined by a two-way ANOVA multiple comparisons test.

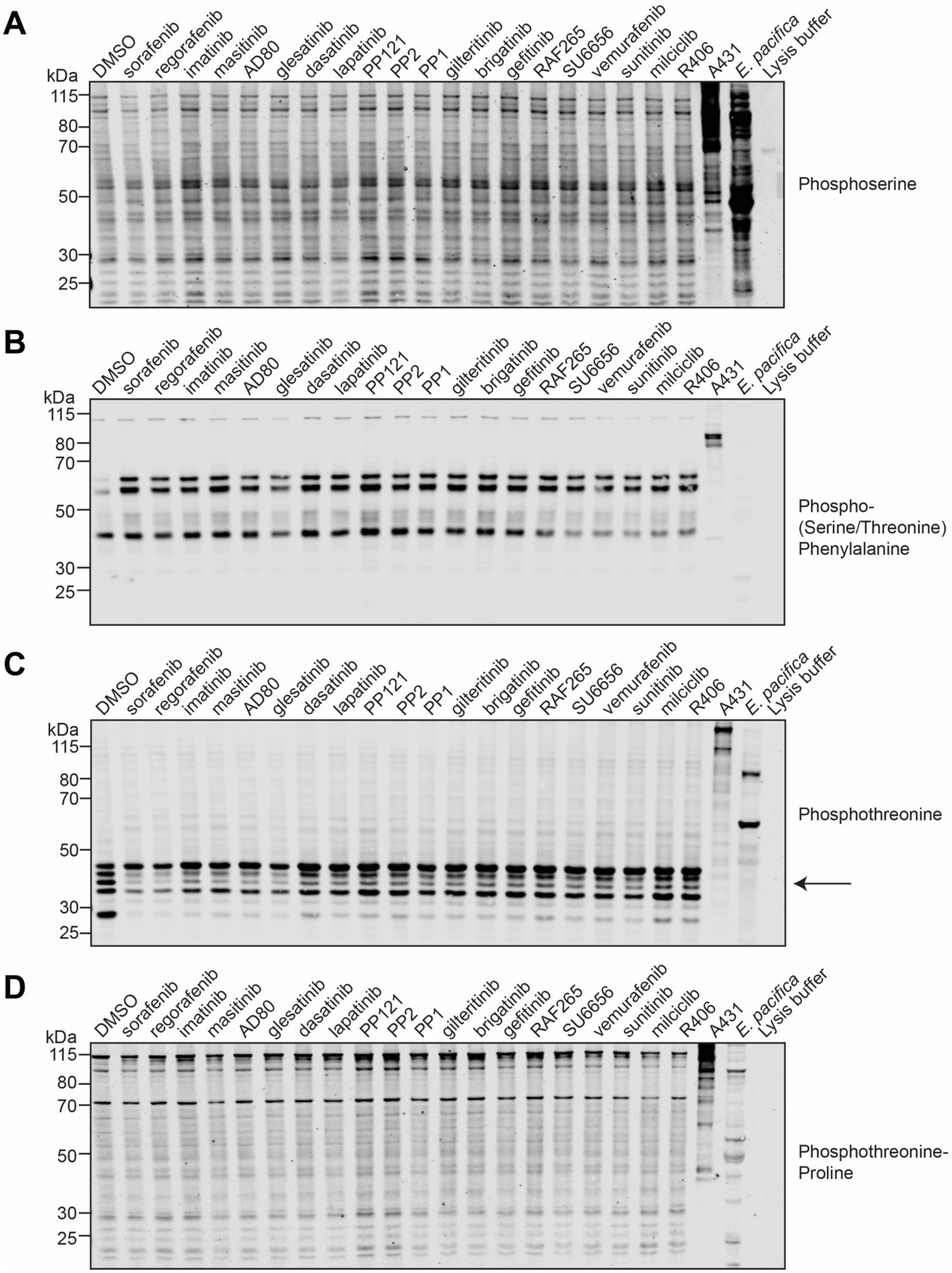
A
DMSO sorafenib regorafenib imatinib masitinib AD80 glesatinib dasatinib lapatinib PP121 PP2 PP1 gilteritinib brigatinib gefitinib RAF265 SU6656 vemurafenib sunitinib milciclib R406 A431 E. pacifica Lysis buffer
kDa
115
80
70
50
30
25
Phosphoserine
B
DMSO sorafenib regorafenib imatinib masitinib AD80 glesatinib dasatinib lapatinib PP121 PP2 PP1 gilteritinib brigatinib gefitinib RAF265 SU6656 vemurafenib sunitinib milciclib R406 A431 E. pacifica Lysis buffer
kDa
115
80
70
50
30
25
Phospho-
(Serine/Threonine)
Phenylalanine
C
DMSO sorafenib regorafenib imatinib masitinib AD80 glesatinib dasatinib lapatinib PP121 PP2 PP1 gilteritinib brigatinib gefitinib RAF265 SU6656 vemurafenib sunitinib milciclib R406 A431 E. pacifica Lysis buffer
kDa
115
80
70
50
30
25
Phosphothreonine
D
DMSO sorafenib regorafenib imatinib masitinib AD80 glesatinib dasatinib lapatinib PP121 PP2 PP1 gilteritinib brigatinib gefitinib RAF265 SU6656 vemurafenib sunitinib milciclib R406 A431 E. pacifica Lysis buffer
kDa
115
80
70
50
30
25
Phosphothreonine-
Proline

**Figure S7.** ***S. rosetta*** **cultures treated with kinase inhibitors did not show a reduction in phosphoserine and phosphothreonine with available antibodies.** *S. rosetta* cultures were treated with 1 μM of the indicated kinase inhibitors for 15 minutes and assessed for changes in phosphoserine and phosphothreonine by four available antibodies. Staining by phosphotyrosine was also conducted and results for tyrosine kinase inhibitors are shown in Figure 3.

(A) Staining with anti-phosphoserine Rb X (EMD Catalog #AB1603) showed no distinct differences in staining with inhibitor treatment.

(B) Staining with anti-phospho(Ser/Thr)-Phe (Cell Signaling Catalog #9631) showed no distinct differences in staining with inhibitor treatment.

(C) Staining with anti-phosphotheronine (Cell Signaling Catalog #9381) showed minor differences in staining at proteins ~40kda with inhibitor treatment.

(D) Staining with anti-phosphothreonine-proline (Cell Signaling Catalog #9391) showed no distinct differences in staining with inhibitor treatment.

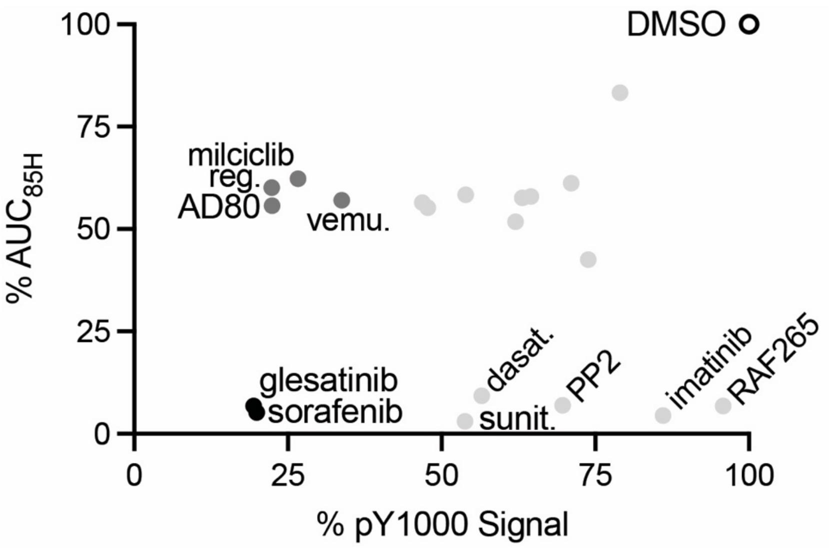


**Figure S8. Sorafenib inhibits *S. rosetta* cell proliferation and tyrosine phosphorylation similarly to glesatinib.** For each of the multi-kinase inhibitors tested (Figure 2, Figure S6-S7), the percent inhibition of phosphotyrosine signal (as detected with the pY1000 antibody, %pY1000) normalized to total protein stain (Figure S6A, Figure 2C) was plotted vs the percentage of the area under the curve over an 85-hour (% $AUC_{85}$) treatment (Figure S6B-D, Figure 2A) of *S. rosetta* cells. Sorafenib and glesatinib most strongly inhibited global phosphotyrosine and cell proliferation. AD80, regorafenib (reg.), milciclib and vemurafenib (vemu.) were strong inhibitors of global phosphotyrosine signaling but moderate inhibitors of *S. rosetta* proliferation. Sunitinib (sunit), dasatinib (dasat.) PP2, and imatinib were strong inhibitors of *S. rosetta* cell proliferation but only moderately or did not decrease the phosphotyrosine signal. Lastly, gilteritinib, lapatinib, PP121, brigatinib, R406, PP1, gefitinib, RAF265 and SU6656 showed only moderate or no effect on *S. rosetta* cell proliferation and phosphotyrosine signaling.

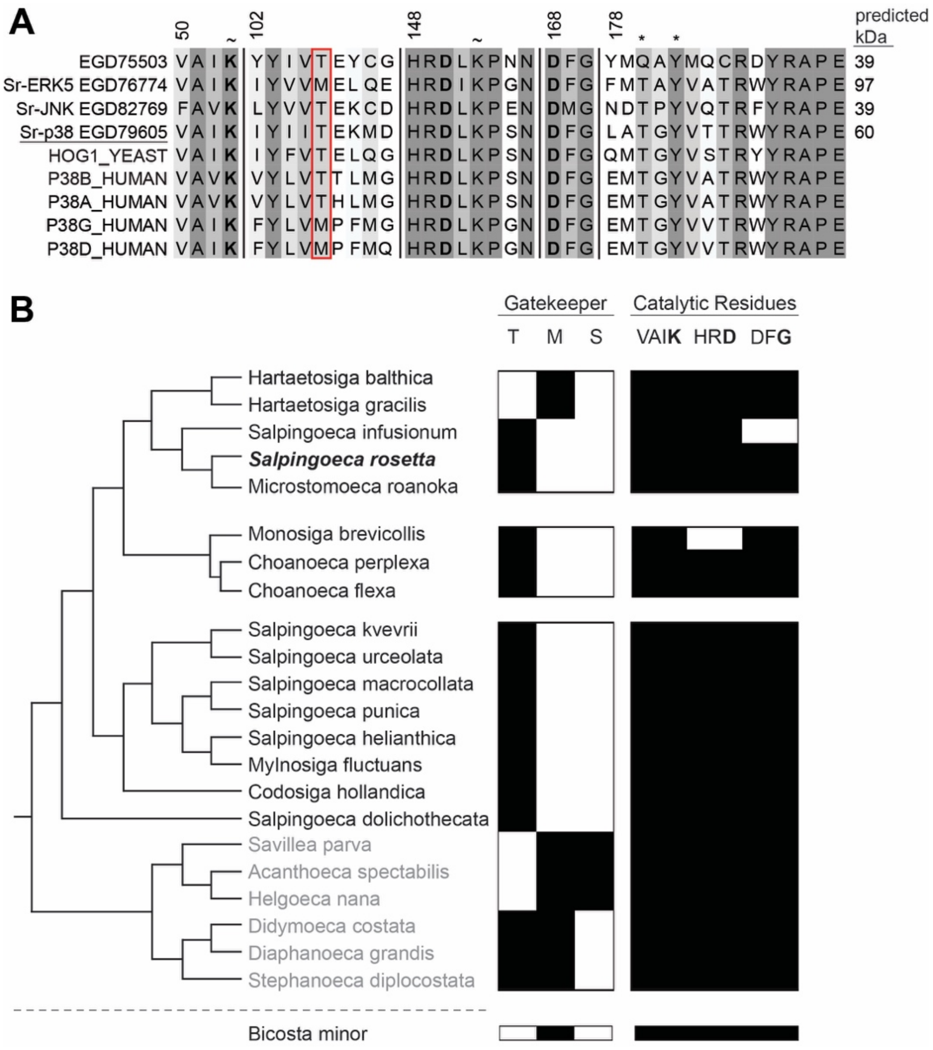


**Figure S9. Choanoflagellates have predicted stress-responsive kinases, including p38 with conserved kinase domain residues**

(A) *S. rosetta*, yeast and human p38 kinases have conserved features. We generated a sequence alignment of the predicted *S. rosetta* stress-responsive kinases [1,7] with Hog1, the p38 homolog from yeast, and the four human p38 kinases. The focus of this study, *Sr*-p38, is underlined. Amino acid positions are referenced to human p38 alpha and are annotated based on predicted function: residues necessary for kinase activity are bolded; (~) indicates lysine residues predicted for ActivX probe binding; (*) indicates sites recognized by the phospho-p38 antibody; the gatekeeper residue position is outlined by a red box.

(B) A choanoflagellate species tree with kinase domain features of representative p38 kinases from available choanoflagellate genomes and transcriptomes[1,8,9]. All choanoflagellates in this tree are predicted to have at least one p38 kinase. Threonine and methionine gatekeepers are predicted in craspedid (black) and acanthoecid (grey) choanoflagellates. *Salpingoeca rosetta* is bolded and italicized.

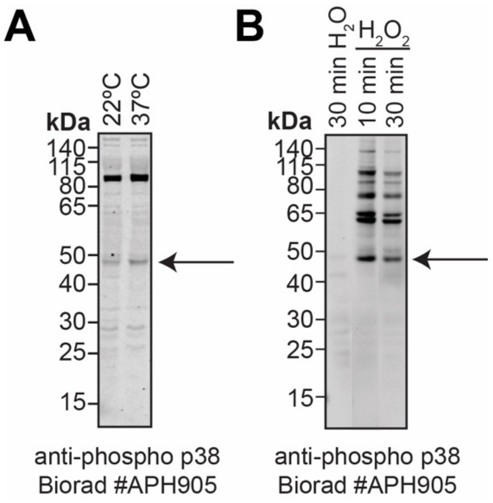


**Figure S10. A second phospho-specific p38 antibody recognizes a band of similar molecular weight.** *S. rosetta* cells were incubated at 37°C or treated with hydrogen peroxide to induce heat shock or oxidative stress. Lysates from the treated cultures were analyzed by western blot and probed with an alternative phospho-specific anti-p38 antibody (p38 MAPK pThr180/pTyr182, Biorad #APH905) to identify if any changes in *Sr*-p38 phosphorylation occurred.

(A) 30 minutes of heat shock at 37°C was sufficient to induce a modest (20%) increase in *Sr*-p38 phosphorylation relative to that seen in cells grown at 22°C.

(B) 10 min or 30 min of treatment with 0.5M $H_2O_2$ at 22°C resulted in a dramatic increase in p38 phosphorylation over that observed in cells treated with water.

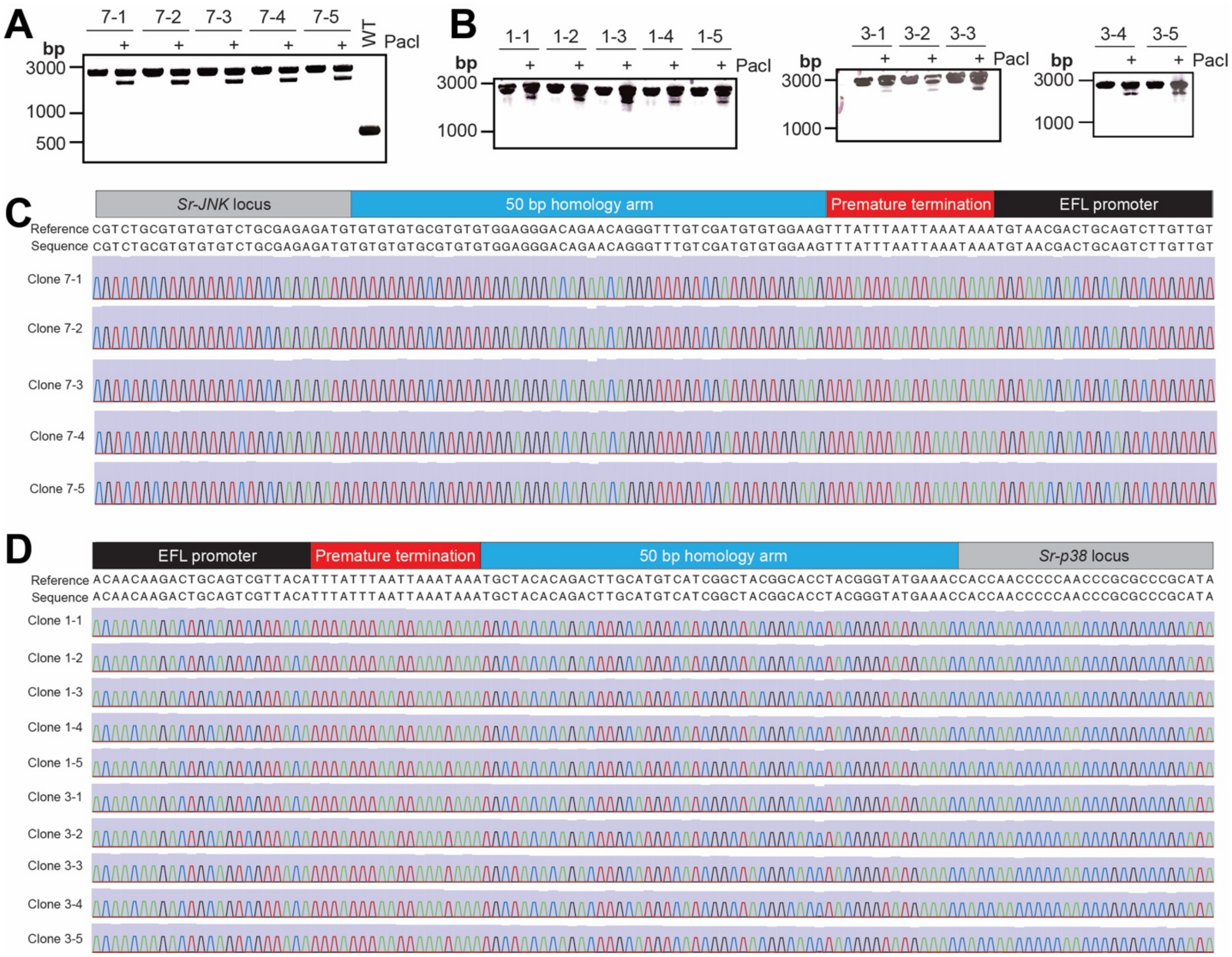


**Figure S11.** ***Sr-p38$^{1\text{-}15}$*** **and** ***Sr-JNK$^{1\text{-}15}$*** **genotyping**

(A) Genotyping PCR of the *Sr-JNK* locus confirms insertion of the premature termination sequence and puromycin resistance cassette in five clones. The approximately 3kb product was digested by PacI, confirming the presence of the premature termination sequence.

(B) Genotyping PCR of the *Sr-p38* locus confirms insertion of the premature termination sequence and puromycin resistance cassette in ten clones. The approximately 3kb product was digested by PacI, confirming the presence of the premature termination sequence.

(C) Nanopore sequencing of *Sr-JNK* PCR products confirms insertion of the premature termination sequence and puromycin resistance cassette in five clones.

(D) Nanopore sequencing of *Sr-p38* PCR products confirms insertion of the premature termination sequence and puromycin resistance cassette in five clones.

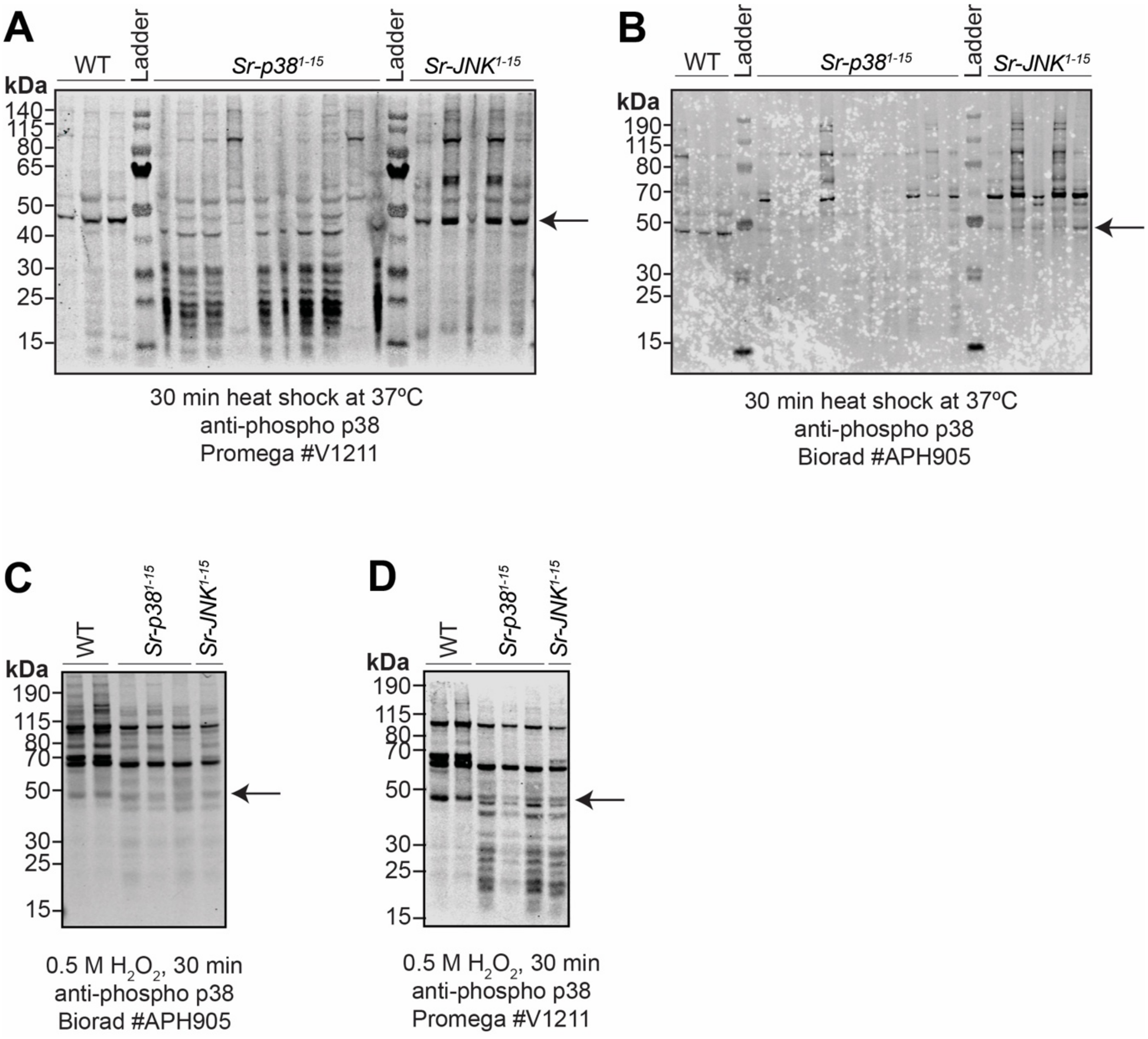


**Figure S12. *Sr-p38*$^{1-15}$ clones lose the phospho-p38 band after heat shock but not after hydrogen peroxide treatment.**

(A) The imaged western blot for Figure 4B that was probed with Anti-ACTIVE® p38 antibody (Promega #V1211) and analyzed by densitometry to quantify differences in *Sr*-p38 phosphorylation. The band at ~45kda (indicated by arrow) in each lane was quantified by densitometry. We attribute antibody denaturing over time to the increase in background on this western blot, compared to Figures 4B and 4C, and could not confirm this supposition because the antibody is no longer commercially available.

(B) The imaged western blot for Figure 4C that was probed with p38 MAPK pThr180/pTyr182 (Biorad #AHP905) and analyzed by densitometry to quantify differences in *Sr*-p38 phosphorylation. The band at ~45kda (indicated by arrow) in each lane was quantified by densitometry.

(C) The phospho-p38 signal induced by hydrogen peroxide in wild-type cells is not decreased in *Sr-p38*$^{1-15}$ knockout cell lines or *Sr-JNK*$^{1-15}$ knockout lines. Two biological replicates of wild-type cells, three of *Sr-p38*$^{1-15}$ and one clone of *Sr-JNK*$^{1-15}$ strains were incubated with 0.5 M hydrogen peroxide for 30 minutes. Lysates from the treated cultures were analyzed by western blot and probed with p38 MAPK pThr180/pTyr182 (Biorad #AHP905).

(D) The phospho-p38 signal induced by hydrogen peroxide in wild-type cells is also not decreased in lysates from *Sr-p38*$^{1\text{-}15}$ knockout cell lines or *Sr-JNK*$^{1\text{-}15}$ knockout lines treated with 0.5 M hydrogen peroxide for 30 minutes and probed with with Anti-ACTIVE® p38 antibody (Promega #V1211).

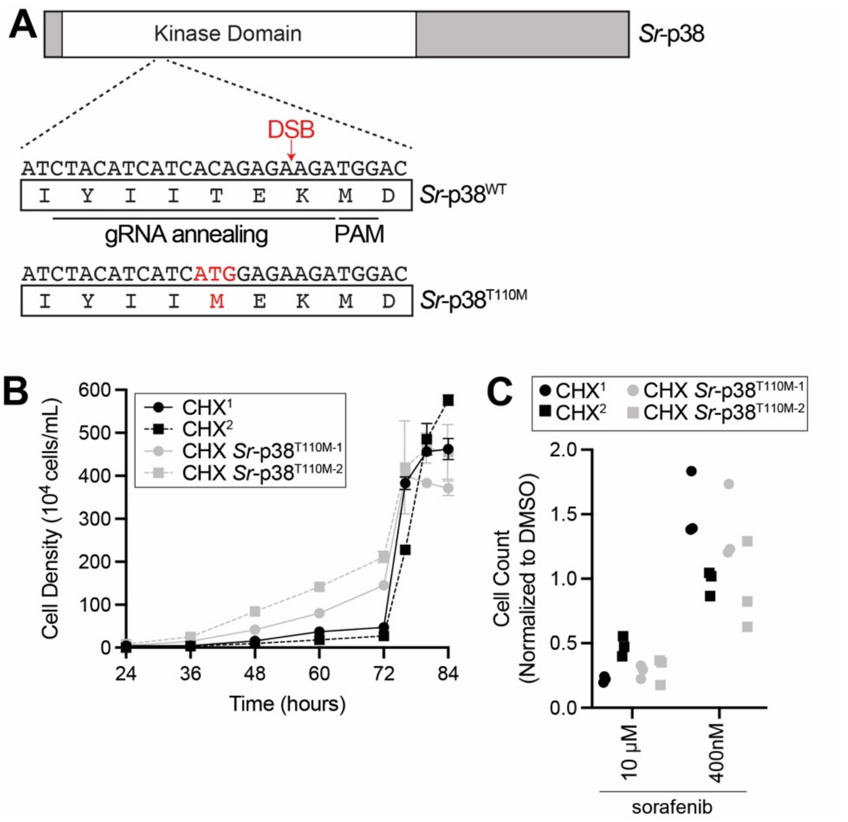


**Figure S13. Inhibitors of human p38 kinases disrupt *S. rosetta* cell proliferation through multiple kinase targets.**

(A) Strategy for generating Sr-p38$^{T110M}$ cell lines. In a previous study, two of four human p38 kinase paralogs with threonine gatekeepers, p38$\alpha$ and p38$\beta$ bound less to skepinone-L, a p38-selective human kinase inhibitor, and sorafenib when the p38 threonine gatekeeper was mutated to methionine. The *S. rosetta* p38 locus was targeted by a guide RNA complexed with Cas9 that anneals near the “gatekeeper” amino acid codon 110 in the kinase domain and directs Cas9 to introduce a double-strand break (DSB) downstream of codon 110. The Cas9-guide RNA complex was coupled with a homology-directed repair template to insert a cassette that encodes a threonine to methionine mutation at position 110. In humans, this mutation desensitizes p38 to sorafenib, and skepinone-L [39]. Co-editing for cycloheximide (CHX) resistance was used to select for clones that were nucleofected with the Cas9 ribonucleoprotein and grew in the presence of CHX, which is resistant to wild-type *S. rosetta* cells. Protein diagrams was created with IBS 2.0 [11].

(B) Two independent clones maintained with CHX resistance, CHX Sr-p38$^{T110M-1}$, and CHX Sr-p38$^{T110M-2}$, do not show a growth defect when compared to CHX$^{1}$ and CHX$^{2}$, two strains that are only cycloheximide-resistant. Data shown are mean cell density with standard deviation. Three technical replicates were conducted per strain, and significance was determined by a two-way ANOVA multiple comparisons test.

(C) CHX Sr-p38$^{T110M}$ strains do not show decreased sensitivity to sorfenib. Two independent CHX Sr-p38$^{T110M}$ and CHX strains were treated with sorafenib and cells were counted after 24 hours. Normalized counts did not differ between the treated CHX Sr-p38$^{T110M}$ and CHX strains. Three technical replicates were conducted per strain, and significance was determined by a two-way ANOVA multiple comparisons test.

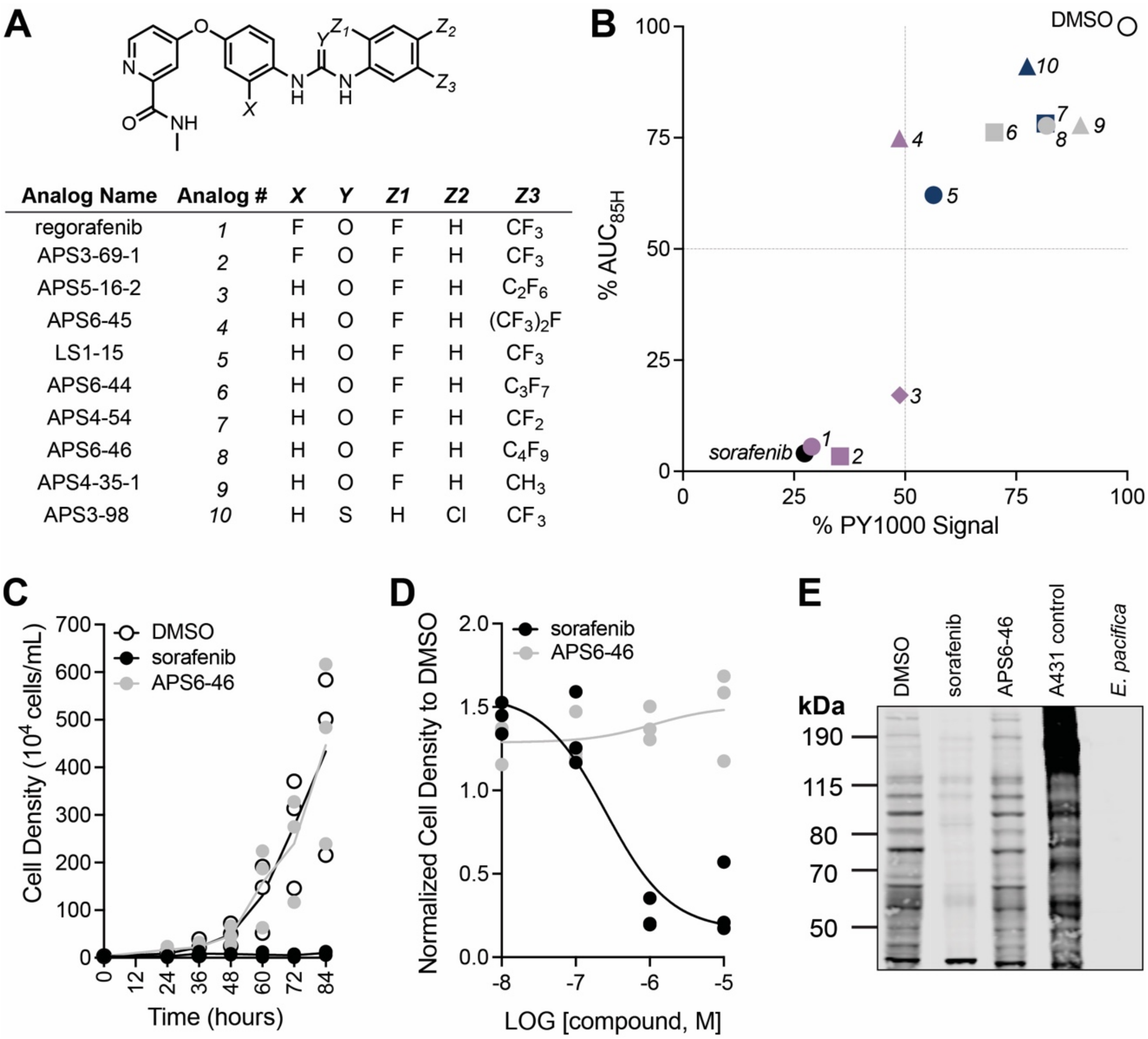


| Analog Name | Analog # | X | Y | Z1 | Z2 | Z3 |
|---|---|---|---|---|---|---|
| regorafenib | 1 | F | O | F | H | $CF_3$ |
| APS3-69-1 | 2 | F | O | F | H | $CF_3$ |
| APS5-16-2 | 3 | H | O | F | H | $C_2F_6$ |
| APS6-45 | 4 | H | O | F | H | $(CF_3)_2F$ |
| LS1-15 | 5 | H | O | F | H | $CF_3$ |
| APS6-44 | 6 | H | O | F | H | $C_3F_7$ |
| APS4-54 | 7 | H | O | F | H | $CF_2$ |
| APS6-46 | 8 | H | O | F | H | $C_4F_9$ |
| APS4-35-1 | 9 | H | O | F | H | $CH_3$ |
| APS3-98 | 10 | H | S | H | Cl | $CF_3$ |

**Figure S14. Sorafenib analogs with increased cap size or decreased cap electronegativity were less potent inhibitors of *S. rosetta* cell proliferation and tyrosine phosphosignaling.**

(A) The chemical structures of additional sorafenib analogs that were tested.

(B) Analogs of sorafenib demonstrated the importance of cap size and substituent electronegativity for compound activity. Other than APS3-69-1 which has a similar cap size to sorafenib, all other analogs tested were less potent inhibitors of *S. rosetta* cell proliferation and/or tyrosine phosphosignaling. The exception was LS1-15, which also has a similar cap size but maintains a similar pattern of X=F being more potent (as in APS3-69-1) that was also observed when comparing sorafenib to regorafenib (Figure 4). The percent inhibition of phosphotyrosine signal (pY1000 antibody, %pY1000) normalized to total protein stain was plotted vs the percentage of the area under the curve over an 85-hour (% $AUC_{85}$) treatment for each analog. Analogs are labeled as analog # for clarity.

(C) *S. rosetta* cultures treated with 1 µM sorafenib in 24-well plates, showed reduced cell density after 48 hours in comparison to APS6-46 and vehicle (DMSO) control. Three biological replicates were conducted per treatment, and each point represents the mean of three measurements from each biological replicate. Normalized cell densities after 48 hours were determined to be reduced if differences between treatments and vehicle (DMSO) were significant (p-value <0.1). Significance was determined by a two-way ANOVA multiple comparisons test.

(D) Inhibition of *S. rosetta* cell proliferation with sorafenib was dose-responsive. Dose-response data are from triplicate treatments of *S. rosetta* cultures treated with sorafenib and APS6-46 for 24 hours in 96-well plates at four different concentrations. The dose-response curve generated is a nonlinear, three-parameter regression with a best-fit pIC50 value of 6.6 for sorafenib.

(E) *S. rosetta* cultures treated with 1 μM sorafenib and regorafenib for 2 hours showed reduced global phosphotyrosine staining whereas cultures treated with APS6-46 did not. EGF-stimulated A431 lysate was used as a positive control for anti-phosphotyrosine (pY1000), and lysate from *E. pacifica*, a bacterial prey of *S. rosetta* was used to control for minor *E. pacifica* protein staining with the pY1000 antibody.

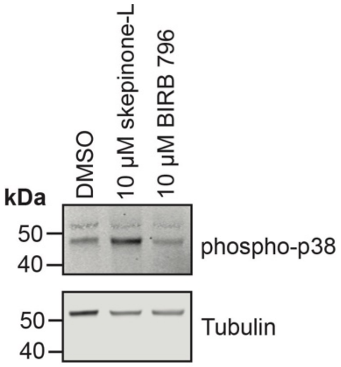


**Figure S15. Skepinone-L and BIRB 796 do not decrease *Sr*-p38 phosphorylation.** *S. rosetta* cultures pretreated with 10 µM of skepinone-L and BIRB 796 for 30 minutes, followed by 30 minutes of heat shock at 37°C were not different from vehicle (DMSO) control. Lysates from the treated cultures were analyzed by western blot with a p38 antibody specific for phosphorylated p38 kinase (phospho-p38) to identify if any changes in *Sr*-p38 phosphorylation occurred.

### Supplementary Table Titles

**Table S1 (separate file).** Normalized *S. rosetta* cell counts by flow cytometry and imaging from *S. rosetta* cells treated with compounds in the screening library and characteristics of those compounds (e.g. kinase targets, inhibitor type, chemical properties)

**Table S2 (separate file).** *S. rosetta* proteins enriched with ActivX probes in the presence or absence of sorafenib and identified with 95% peptide and protein confidence score and at least two unique peptides.

### Supplementary Movies

**Movie S1 (separate file)**. Two-hour 40 minute timecourse of *S. rosetta* cells treated with 10 μM glesatinib. Arrows point to three cells that undergo cell lysis.

**Movie S2 (separate file)**. One-hour timecourse of *S. rosetta* cells treated with 10 μM sorafenib. Most cells elongate and extend filopodia (see Movie S2 for zoomed in video)

**Movie S3 (separate file)**. Cropped one-hour timecourse of an *S. rosetta* cell treated with 10 μM sorafenib (cropped from Movie S1). Arrow points to area of cell body that elongates during treatment.

**Movie S4 (separate file)**. Three-hour 30 minute timecourse of *S. rosetta* cells treated with vehicle (DMSO) control. Arrows point to cells that undergo cell division.
